# Allelic compatibility in plant immune receptors facilitates engineering of new effector recognition specificities

**DOI:** 10.1101/2022.10.10.511592

**Authors:** Adam R. Bentham, Juan Carlos De la Concepcion, Javier Vega Benjumea, Sally Jones, Melanie Mendel, Jack Stubbs, Clare E. M. Stevenson, Josephine H.R. Maidment, Mark Youles, Jiorgos Kourelis, Rafał Zdrzałek, Sophien Kamoun, Mark J. Banfield

## Abstract

Engineering expanded effector recognition in plant immune receptors is a promising prospect for generating new disease resistant crop varieties. However, modification of plant NLR receptors has proven challenging due to the lack of understanding of their context as part of complex immune systems. Here, we demonstrate a new avenue for NLR-mediated engineering that exploits the allelic diversity in the Pik NLR pair to allow for the generation of receptors with expanded recognition specificities, which would otherwise result in constitutive cell death. This work lays the foundation for the incorporation of new effector recognition motifs into the Pik system and advances the development of designer NLRs that can be tailored to specific secreted pathogen signatures.

**Abstract:** Engineering the plant immune system offers genetic solutions to mitigate crop diseases caused by diverse agriculturally significant pathogens and pests. Modification of intracellular plant immune receptors of the nucleotide-binding leucine rich repeat (NLRs) superfamily for expanded recognition of pathogen virulence proteins (effectors) is a promising approach for engineering novel disease resistance. However, engineering can cause NLR autoactivation, resulting in constitutive defence responses that are deleterious to the plant. This may be due to plant NLRs associating in highly complex signalling networks that co-evolve together, and changes through breeding or genetic modification can generate incompatible combinations, resulting in autoimmune phenotypes. We have previously shown how alleles of the rice NLR pair Pik have differentially co-evolved, and how sensor/helper mismatching between non-co-evolved alleles triggers constitutive activation and cell death (De la Concepcion et al., 2021b). Here, we dissect incompatibility determinants in the Pik pair and found that HMA domains integrated in Pik-1 not only evolved to bind pathogen effectors but also likely co-evolved with other NLR domains to maintain immune homeostasis. This explains why changes in integrated domains can lead to autoactivation. We then used this knowledge to facilitate engineering of new effector recognition specificities overcoming initial autoimmune penalties. We show that by mismatching alleles of the rice sensor and helper NLRs Pik-1 and Pik-2, we can enable the integration of synthetic HMA domains with novel and enhanced recognition of an effector from the rice blast fungus. Taken together, our results reveal a new strategy for engineering NLRs, which has the potential to allow an expanded set of integrations and therefore new disease resistance specificities in plants.

## Introduction

Engineering the plant immune system is a promising genetic solution to prevent pathogen infection thereby reducing crop losses and global food insecurity caused by plant pathogens (Bentham et al., 2020; Outram et al., 2022). Nucleotide-binding leucine rich repeat receptors (NLRs) are intracellular immune proteins that trigger robust defence responses upon recognition of pathogen virulence proteins (effectors) delivered into the host during infection (Burdett et al., 2019; Jones et al., 2016). Effector recognition by NLRs often culminates in cell death that isolates the invading pathogen and confers resistance (Jones et al., 2016; Maruta et al., 2022). Due to their effective and specific responses to plant pathogens, engineering of NLRs to increase their effector recognition specificities is a promising approach to boost disease resistance (Marchal et al., 2022; Monteiro and Nishimura, 2018; Outram et al., 2022).

NLRs are typically modular tripartite proteins that consist of an N-terminal signalling domain, either a coiled-coil (CC) domain, a CC domain with homology to RPW8 (CC_R_) or Toll-interleukin-1 receptor (TIR) domain, a central nucleotide-binding (NB) domain, and a C-terminal leucine rich repeat (LRR) domain (Jones et al., 2016; Lüdke et al., 2022; Takken and Goverse, 2012). NLR proteins can function as singletons, pairs and in networks, and utilize several mechanisms to detect and respond to pathogen effectors (Adachi et al., 2019; Cesari, 2018; Wu et al., 2018). One such mechanism is the use of integrated domains, which function as effector baits embedded within the canonical NLR architecture (Baggs et al., 2017; Cesari et al., 2014). Integrated domains often share homology to pathogen host targets and effector binding results in NLR activation (Białas et al., 2018; Cesari, 2018; Kroj et al., 2016; Sarris et al., 2016). Due to their role in effector recognition, integrated domains are key targets for engineering disease resistance in NLRs and only a few mutations to these domains can lead to novel effector recognition profiles (Cesari et al., 2022; De la Concepcion et al., 2019; Liu et al., 2021; Maidment et al., 2022; Zhang et al., 2022).

Most characterised NLRs that contain integrated domains (NLR-IDs) are found as a part of a sensor/helper receptor pair, where the NLR-ID is referred to as the sensor, and its signalling partner, the helper (Adachi et al., 2019; Cesari, 2018; Feehan et al., 2020). Some of the best characterised examples of paired NLRs are the Arabidopsis RRS1/RPS4 pair which encodes an RRS1-integrated WRKY domain (Le Roux et al., 2015; Mukhi et al., 2021; Zhang et al., 2017) and the rice NLR pairs RGA5/RGA4 and Pik-1/Pik-2 (Cesari et al., 2013; Kanzaki et al., 2012; Zdrzałek et al., 2020). The RGA5/RGA4 and Pik pairs harbour an integrated HMA domain in their sensor NLRs RGA5 and Pik-1 that directly bind and recognise MAX (*M. oryzae* avirulence and ToxB-like) effectors (Guo et al., 2018; Maqbool et al., 2015). While both RGA5 and Pik-1 contain an integrated HMA domain, their domain architecture is distinct with the Pik-1 HMA domain located between the CC and NB domains, and the RGA5 HMA domain located at the C-terminus, after the LRR (Cesari et al., 2013; Kanzaki et al., 2012; Maqbool et al., 2015; Ortiz et al., 2017). Further, these HMA domains provide distinct effector recognition specificities for AVR-Pik, AVR-Mgk1, and AVR-Pia/AVR1-CO39 effectors respectively (Białas et al., 2018; Sugihara et al., 2022).

The HMA domains of Pik-1 and RGA5 use spatially distinct protein interfaces for effector recognition (De la Concepcion et al., 2021a, 2018; Guo et al., 2018; Varden et al., 2019). Recent studies have reported HMA domain engineering to be an effective way to generate new resistance specificities for rice against the rice blast pathogen *M. oryzae*. In particular, three separate studies have shown the RGA5 HMA domain can be engineered to recognise other MAX effectors. One study showed engineered resistance to *M. oryzae* isolates carrying AVR-Pib in rice (Liu et al., 2021), and two studies engineered recognition of AVR-Pik in *N. benthamiana*, one of which was able to provide resistance in rice (Cesari et al., 2022; Zhang et al., 2022). Extensive study of the Pik-1 HMA domain has also demonstrated this HMA domain to be amenable to engineering (De la Concepcion et al., 2021a, 2019; Maidment et al., 2022). Recently, it has been shown the Pik-1 HMA can be substituted for VHH nanobody fusions that act as synthetic effector recognition domains, demonstrating the flexibility of the Pik system for mutation or substitution of new integrated domains (Kourelis et al., 2021).

Despite some success, plant immune receptor engineering remains challenging. NLRs exist in complex, regulated systems and as a consequence some changes in NLRs that expand recognition can also result in constitutive defence signalling that is deleterious for plant growth (Białas et al., 2021; Maidment et al., 2022; Tamborski et al., 2022). NLR-mediated autoimmunity has been well documented, with hybrid necrosis phenotypes as a result of crosses linked to incompatible pairing of NLRs (Bomblies et al., 2007; Chae et al., 2014; Kourelis and Adachi, 2022; Tran et al., 2017), presenting a bottleneck to producing new resistant crop varieties either by breeding or precision protein engineering.

Recently, we demonstrated alleles of the rice Pik NLR pair have differentially co-evolved, likely driven by their differences in recognition specificity for *Magnaporthe oryzae* AVR-Pik effector variants (De la Concepcion et al., 2021b). The Pik alleles Pikp and Pikm have undergone functional diversification, with multiple changes in their integrated HMA domain that result in different recognition specificities for AVR-Pik variants (Bialas et al., 2021; De la Concepcion et al., 2021). Where the Pikp-1 sensor is restricted to detecting AVR-PikD, Pikm-1 is able to recognise AVR-PikD, AVR-PikE and AVR-PikA (De la Concepcion et al., 2018; Kanzaki et al., 2012). The helper NLRs Pikp-2 and Pikm-2 also appear to have undergone diversification to match their sensor partners that results in a one-way incompatibility between Pik alleles (De la Concepcion et al., 2021b). While Pikp-2 can be used as a helper with Pikm-1 to recognise AVR-Pik, the Pikp-1/Pikm-2 combination results in constitutive cell death in *N. benthamiana*. This incompatibility between Pikp-1 and Pikm-2 is linked to a single polymorphism in the NB-ARC of Pikm-2, which when mutated to the equivalent residue of Pikp-2 reinstates compatibility (De la Concepcion et al., 2021b).

Here, we demonstrate the autoactivity triggered by the engineering of Pikm-1 for expanded effector recognition capabilities can be attenuated by the co-expression with Pikp-2 without compromising receptor function. For this, we delineate the basis for receptor incompatibility between Pikp and Pikm alleles, describing Pikp-2 as a facilitator for integration of new integrated domains into Pikm-1. By mismatching Pikm-1 with Pikp-2, we can integrate the RGA5 HMA domain into Pikm-1, enabling further engineering of RGA5 HMA to recognise multiple AVR-Pik variants. Finally, we structurally and biophysically characterise the interaction between a synthetic AVR-Pik-binding mutant of RGA5 HMA and AVR-Pik effectors, highlighting the importance of binding affinity between effector and bait for immune recognition. These results emphasize the importance of tuning receptor pairs in engineering and supplies a novel approach to NLR engineering that will aid in the implementation of modified immune receptors with expanded effector recognition specificities outside of that previously observed in nature.

## Results

### The Pik-HMA domain is not required for effector-independent immune signalling

Mismatched pairing of the Pikp-1/Pikm-2 alleles triggers constitutive cell death in the absence of an effector binding to the integrated HMA domain (De la Concepcion et al., 2021b). To better assess the role of the Pik HMA domain in signalling, activation and autoimmunity outside of effector binding, we used an HMA-absent Pikp-1 variant (Pikp-1^ΔHMA^) where the HMA domain of Pikp-1 was substituted with the unrelated NOI domain from the rice NLR Pii-2 (Pii-2 residues Glu1016 to Lys1052) (Fujisaki et al., 2017). Constitutive cell death was observed upon co-expression of Pikp-1^ΔHMA^ with Pikm-2 in *N. benthamiana*. However, like wildtype Pikp-1, co-expression of Pikp-1^ΔHMA^ with Pikp-2 did not result in cell death (**Figure 1** – **Appendix 1A**). Then we found autoactivity induced by co-expression of Pikp-1^ΔHMA^/Pikm-2 is determined by the Pik-2 Asp230Glu polymorphism (Pikp-2^D230E^/Pikm-2^E230D^) (**Figure 1** – **Appendix 1A**) as described for wildtype Pik NLRs (De la Concepcion et al., 2021b). An Asp230Glu mutation into Pikp-2 resulted in constitutive activation when co-expressed with Pikp-1^ΔHMA^, and the reciprocal mutation, Glu230Asp, in Pikm-2 abolished autoactivity (**Figure 1** – **Appendix 1A**). This suggests the integrated HMA domain acts as an effector binding domain but is not required for downstream NLR signalling and cell death.

**Figure 1.**
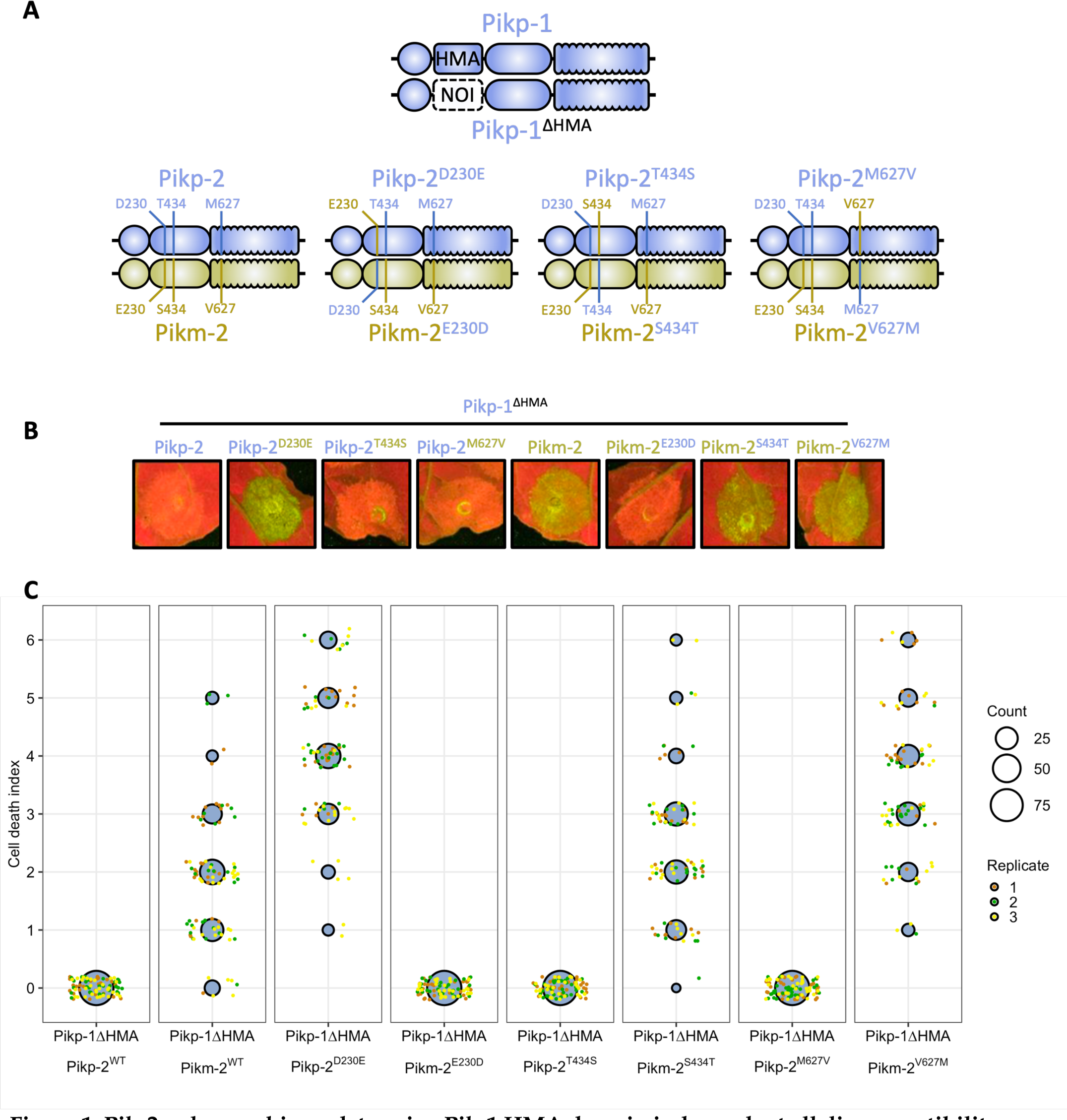
Pik-2 polymorphisms determine Pik-1 HMA domain independent allelic compatibility. **A)** Schematic diagram highlighting the domain architecture of Pikp-1 and Pikp-1^ΔHMA^ sensor NLRs and the polymorphisms between the Pikp-2 and Pikm-2 helper NLRs. **B)** The HMA domain of Pikp-1 is not required for effector-independent cell death in *N. benthamiana*; compatibility remains with Pikp-2 but not Pikm-2. Reciprocal D230E and E230D mutations in Pikp-2 and Pikm-2 flip the compatibility of the helper NLRs for Pikp-1^ΔHMA^. **C)** Cell death scoring for repeats of Pikp-1^ΔHMA^ co-expressed with Pikp-2, Pikm-2, and mutants in *N. benthamiana* represented as dot plots. The total number of repeats was 75 per sample. For each sample, all the data points are represented as dots with a distinct colour for each of the three biological replicates; these dots are jittered around the cell death score for visualisation purposes. The size of the central dot at each cell death value is proportional to the number of replicates of the sample with that score. Quantification and statistical analysis of these results can be found in **Appendix 1 A**.

### Incompatibility between alleles of the Pik NLR pair is linked to regions within the sensor and the helper

As the integrated HMA is not required for immune activation of the Pik pair, and is the most variable domain between the Pikp-1 and Pikm-1 alleles (Costanzo and Jia, 2010), we hypothesised the Pikm-HMA domain co-evolved with Pikm-2 to supress autoactivation mediated by Pik-2 Asp230Glu polymorphism. To test this, we exchanged the integrated HMA domains between sensor alleles Pikp-1 (pHMA) and Pikm-1 (mHMA), to create Pikp-1^mHMA^ and Pikm-1^pHMA^. Pikp-1^mHMA^ and Pikm-1^pHMA^ were co-expressed in *N. benthamiana* with either the Pikp-2 or Pikm-2 helper and challenged with AVR-PikD or mCherry to test for effector activation and autoimmunity, respectively (**Figure 2 A, Figure S1 A** – **Appendix 1 B**). Expression of the Pikm-1^pHMA^ with Pikm-2 resulted in effector-independent cell death; however, this sensor was not autoactive in the presence of Pikp-2 and was able to respond to AVR-PikD. By contrast, Pikp-1^mHMA^ was not autoactive when co-expressed with either the Pikp-2 or Pikm-2 helpers and cooperated with either helper to respond to AVR-PikD. These data demonstrate the integrated domain of the Pik-1 sensor contributes to the compatibility between the Pik sensor and helper NLRs.

**Figure 2.**
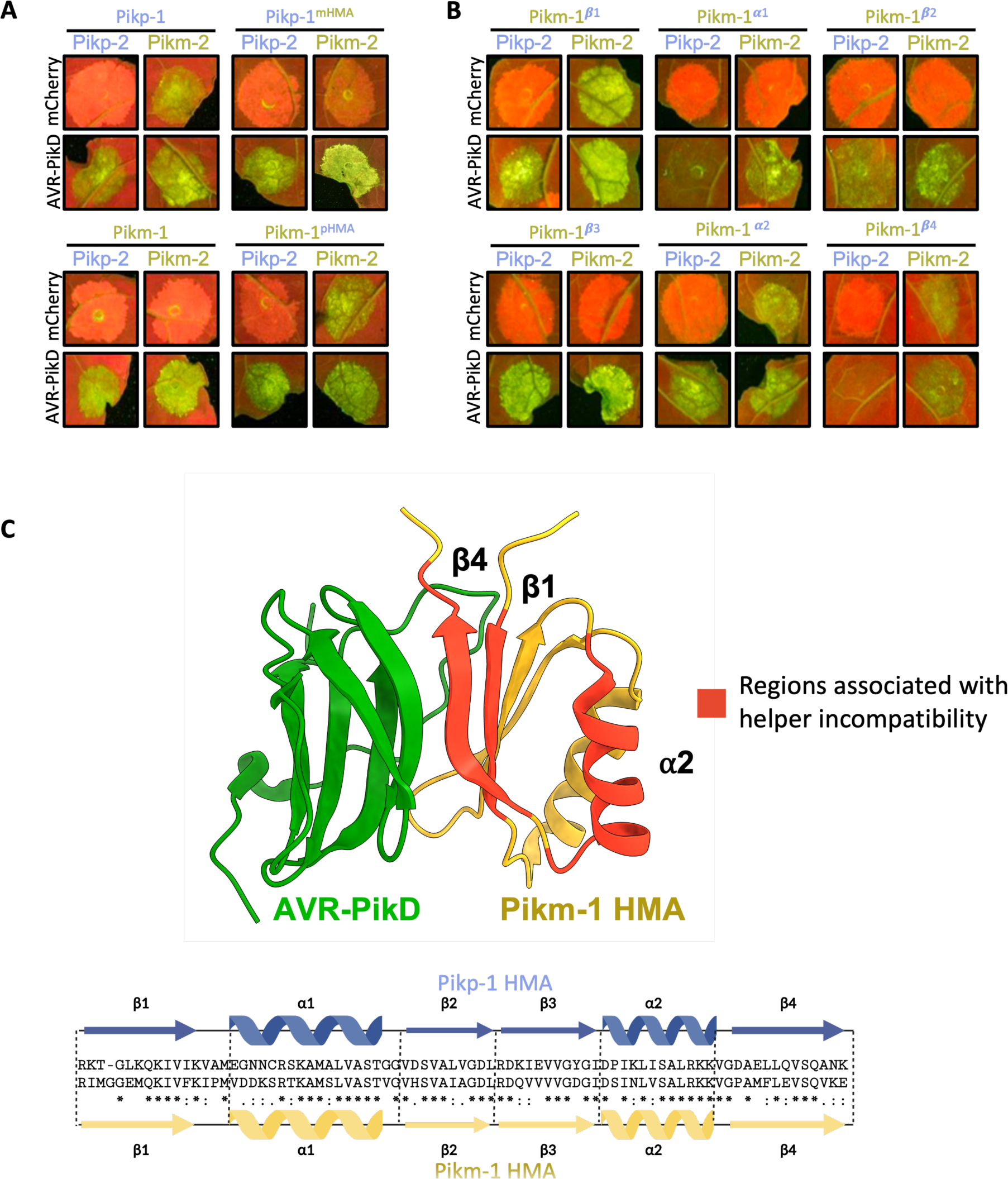
The HMA domain of Pik-1 is important for compatibility with Pik-2 helpers. **A)** Co-expression of Pikp-1 with Pikm-2 triggers effector-independent cell death in *N. benthamiana.* Integration of the Pikm-1 HMA into Pikp-1 facilitates Pikp-1 compatibility with Pikm-2, whereas incorporation of the Pikp-1 HMA into Pikm-1 abolishes compatibility with Pikm-2. Quantification and statistical analysis of these results are shown in **Figure S1 A, Appendix 1 B**. **B)** Incompatibility of the Pikp-1 with Pikm-2 in *N. benthamiana* is linked to the α2 helix, β1, and β4 strands of the HMA domain, with Pikm-1 HMA chimeras carrying the Pikp-1 secondary structure elements resulting in effector-independent cell death when co-expressed with Pikm-2. Quantification and statistical analysis of these results can be found in **Figure S1 B, Appendix 1 C**. **C)** Some regions of the HMA domain that are involved in sensor/helper incompatibility (red) are shared with the AVR-Pik binding interface (PDB ID: 6FU9).

To gain a better understanding of which features of the HMA domain are involved in sensor/helper compatibility, we generated chimeras by introducing secondary structures from the Pikp-1 HMA into the Pikm-1 HMA and tested for autoactivation in the presence of Pikm-2. The Pik HMA maintains a four-strand β-sandwich fold (β1 – β4) flanked by two helices (α1 and α2) (De la Concepcion et al., 2018). For this experiment we generated six chimeric sensors: Pikm-1^β1^, Pikm-1^α1^, Pikm-1^β2^, Pikm-1^β3^, Pikm-1^α2^, and Pikm-1^β4^, and these were co-expressed with Pikm-2 and AVR-PikD or mCherry in *N. benthamiana* (**Figure 2 B, Figure S1 B – Appendix 1 C**). Of the six mutants, only Pikm-1^β1^, Pikm-1^α2^, and Pikm-1^β4^ resulted in effector-independent cell death. However, not all the residues of the β1 and β4 strands make significant contributions to the AVR-Pik binding interface of the HMA (**Figure 2 C**, **Figure S2, Figure S3** – **Appendix 1 D-F**), implying that some residues not directly involved in the binding to the effector can have a regulatory role in NLR activation.

Following the observation that the β1, α2, and β4 HMA secondary structures may be involved in helper incompatibility, we created single point mutations of the polymorphic residues between Pikp and Pikm HMA domains in these secondary structures (**Figure S2, Figure S3 – Appendix 1 D-F**) to assess their individual contributions to sensor/helper compatibility. When co-expressed in *N. benthamiana*, few of the Pikm-1 mutants influenced compatibility with Pikm-2 in contrast to our observations with the α2, β1, and β4 chimeras, which points towards a certain threshold for change in the HMA being tolerated by the system (**Figure S2, Figure S3 – Appendix 1 D-F**). Notable exceptions to this were the deletion of Gly186 in β1 and the Pro252Asp substitution in β4, which resulted in strong autoactivity in the presence of Pikm-2. Why these two mutations result in such strong autoactivity is unclear, but could be related to both causing large-scale structural changes, as the removal of a residue (ΔG186) or mutation of a proline (Pro252) could impact secondary structure formation and affect the ability of the mHMA to prevent autoactivity in the presence of Pikm-2.

Taken together, these data demonstrate elements of the Pik-1 sensor and Pik-2 helper contribute to receptor compatibility, as the HMA domain is not required for cell death signalling but has evolved to accommodate for changes in Pik-2 that would otherwise result in constitutive activation.

### Integration of the RGA5 HMA domain into Pik-1 is facilitated by allelic mismatching

Our results also suggest Pikp-2 may be more accommodating of changes in the Pik-1 sensor than Pikm-2, even tolerating the complete substitution of the integrated HMA by an unrelated domain without inducing autoactivity. We hypothesised the ability of Pikp-2 to accommodate changes in the integrated domain would allow for integration of an HMA domain that would normally result in autoactivity. To test this, we made a chimera of Pikm-1 carrying the HMA domain from the rice NLR RGA5. Using multiple sequence alignment and structural visualisation in ChimeraX (Pettersen et al., 2021), we defined residues 997 – 1071 from the RGA5 HMA to be the identical boundaries of the Pikm-1 HMA domain. It was important to make sure the size of the HMA incorporated into the Pikm-1 chassis was identical to the Pikm-1 HMA due to our previous observation that removal of a single residue from the HMA (ΔG186) resulted in strong autoactivity.

Co-expression of the Pikm-1^RGA5^ chimera with Pikm-2 in *N. benthamiana* resulted in a strong effector-independent cell death response. However, no cell death was observed upon co-expression of Pikm-1^RGA5^ with Pikp-2 (**Figure 3 – Appendix 1 G**). Therefore, the Pikp-2 helper allows the integration of the RGA5 HMA into Pikm-1. To test whether the Pikm-1^RGA5^ chimera is a functional receptor, the Pikm-1^RGA5^/Pikp-2 combination was co-expressed with AVR-Pia. Co-expression of Pikm-1^RGA5^, Pikp-2 and AVR-Pia in *N. benthamiana* resulted in weak cell death, significantly weaker than the cell death elicited by RGA4/RGA5 in response to AVR-Pia (**Figure 3 – Appendix 1 G**), but comparable to the cross-reactivity of Pikp-1/Pikp-2 with AVR-Pia previously observed in *N. benthamiana* (Varden et al., 2019). Furthermore, we sought determine whether the RGA5 HMA integrated into Pikm-1 could complement the function of the Pikm-1 HMA and respond to AVR-PikD (**Figure S4 – Appendix 1 H**). Co-expression of Pikm-1^RGA5^/Pikp-2 with AVR-PikD did not trigger a cell death response in *N. benthamiana*, demonstrating the RGA5 HMA cannot substitute the Pikm-1 HMA as an AVR-Pik recognition module when integrated into Pikm-1.

**Figure 3.**
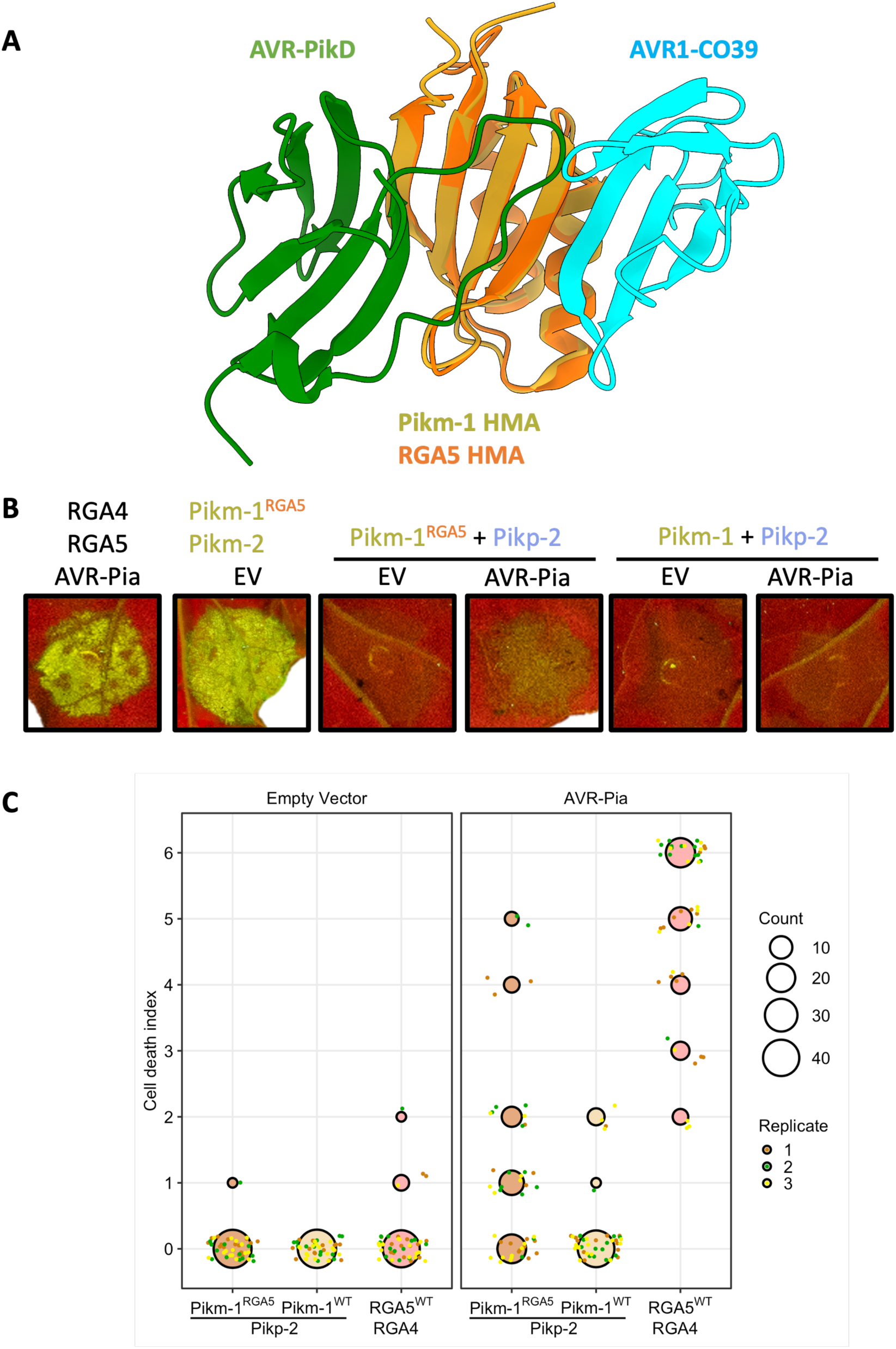
Autoactivity following integration of the RGA5 HMA domain into Pikm-1 is relieved by allelic mismatch with Pikp-2, but only weakly responds to AVR-Pia. **A)** Schematic structural alignment of the RGA5 HMA domain (orange; PDB ID: 5ZNG) with the Pikm-1 HMA domain (gold; PBD ID: 6FU9) showing the different binding interfaces of these HMAs for the AVR-Pik (green) and AVR1-CO39 (cyan)/AVR-Pia (not shown) effectors. B) Co-expression of Pikm-1^RGA5^ with Pikp-2 supresses effector independent cell death and responds weakly to AVR-Pia. C) Cell death scoring of wildtype Pikm-1 and Pikm-1^RGA5^ co-expressed with Pikp-2 in *N. benthamiana* represented as dot plots. The total number of repeats was 45 per sample. For each sample, all the data points are represented as dots with a distinct colour for each of the three biological replicates; these dots are jittered around the cell death score for visualisation purposes. The size of the central dot at each cell death value is proportional to the number of replicates of the sample with that score. Statistical analyses of these results are shown in Appendix 1 G.

These data demonstrate the Pikp-2 helper can be used to facilitate the integration of new domains into the Pikm-1 sensor that would otherwise result in autoactivity/incompatibility when paired with Pikm-2.

### The RGA5 HMA domain can be engineered to recognise AVR-Pik from within the Pik-1 chassis

To test whether the RGA5 HMA can act as an effector recognition module in the Pikm-1 receptor, we engineered AVR-Pik recognition in the RGA5 HMA. Using a host target of AVR-Pik, OsHIPP19 (Maidment et al., 2021) as a structural template for AVR-Pik binding we generated an RGA5 variant, termed the AVR-Pik binding (APB) mutant (**Figure S5**). The APB mutant contains the point mutations Glu1033Asp, Val1039Gln, Met1065Gln, Leu1068Glu, Glu1070Lys, and Lys1071Glu that localise to a potential AVR-Pik binding interface of the RGA5 HMA. Next, we generated a Pikm-1 chimera containing the RGA5-APB mutant HMA (Pikm-1^APB^) and co-expressed it with Pikp-2 and the AVR-Pik variants AVR-PikD, AVR-PikC and AVR-PikF in *N. benthamiana.* The Pikm-1^APB^ chimera was able to trigger cell death in response to AVR-PikD, AVR-PikC and AVR-F (**Figure 4 A, B – Appendix 1 I**). We also tested whether Pikm-1^APB^ could recognise AVR-Pia. However, as for the Pikm-1^RGA5^ chimera, we only observed a weak cell death response (**Figure 4 A, B – Appendix 1 I**).

**Figure 4.**
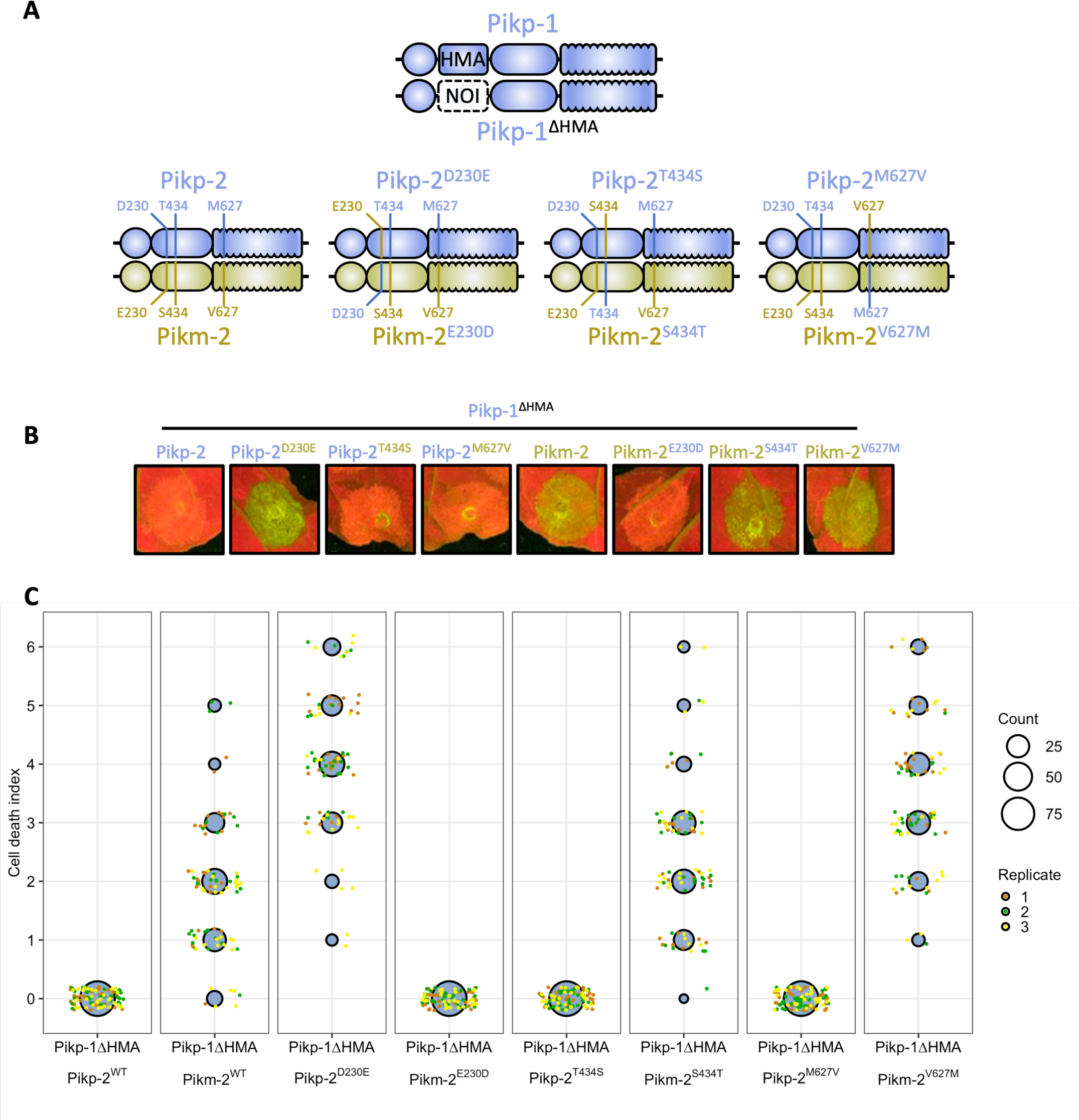
The RGA5 APB mutant binds and recognises AVR-Pik when integrated into Pikm-1. **A)** Pikm-1^APB^ chimera responds to all variants of AVR-Pik tested and activates cell death when co-expressed with Pikp-2 in *N. benthamiana*, but like Pikm-1^RGA5^, it only weakly responds to AVR-Pia. **B)** Cell death scoring of Pikm-1^APB^ co-expressed with AVR-Pik variants D, C and F in *N. benthamiana* represented as dot plots. The total number of repeats was 80 per sample. For each sample, all the data points are represented as dots with a distinct colour for each of the three biological replicates; these dots are jittered around the cell death score for visualisation purposes. The size of the central dot at each cell death value is proportional to the number of replicates of the sample with that score. Statistical analyses of these results are shown in **Appendix 1 I C)** Co-immunoprecipitation of Pikm-1^APB^ with different MAX effectors shows association with AVR-Pik variants, but not AVR-Pia, in planta. Dotted line denotes separate membrane exposures of the same membrane. **D)** Surface plasmon resonance sensograms for the interaction of HMA domains of Pikm-1, RGA5 and RGA5 APB mutant with effectors AVR-PikD, AVR-PikC, AVR-PikF and AVR-Pia. Non-MAX effector AVR-Pii was added as a negative control. Response units for each labelled protein concentration are shown with the residuals plot beneath (SPR acceptance guides as determined by Biacore software are shown as green and red lines in the residuals plots). Concentration of each protein in the assay is indicated next to their corresponding name. Each experiment was repeated a minimum of 3 times, with similar results.

To observe whether cell death correlates with effector binding in planta, we performed co-immunoprecipitation (co-IP) assays with FLAG-tagged Pikm-1^APB^ and 4xMYC tagged AVR-PikD, AVR-PikC, AVR-PikF, AVR-Pia, and PWL2 as negative control. We observed bands corresponding to AVR-PikD, AVR-PikC and AVR-PikF in Pikm-1^APB^ samples after FLAG pulldown, whereas only AVR-PikD was pulled down by wildtype Pikm-1 (**Figure 4 C**). Correlating with the weak response in cell death assays, we were unable to observe association between Pikm-1^APB^ and AVR-Pia. Taken together, these data demonstrate the RGA5 HMA domain can be engineered to respond to AVR-Pik in planta in the context of the Pikm-1 receptor. However, incorporation of the RGA5 HMA in the Pikm-1 chassis is not sufficient for robust recognition of AVR-Pia.

### The affinity of HMA domains for effectors underpins recognition phenotypes in Pik-1 chimeras

We hypothesized a lower affinity of the APB HMA for AVR-Pia compared to AVR-Pik is responsible for the differences in cell death phenotypes. The interaction between AVR-Pia/RGA5-HMA is known to be much weaker when compared with AVR-Pik/Pik-HMA (De la Concepcion et al., 2018; Guo et al., 2018; Ortiz et al., 2017). To investigate this, we performed surface plasmon resonance (SPR) to determine affinities of the RGA5, Pikm-1 and APB HMA domains for different MAX effectors.

We purified AVR-PikD, AVR-PikC, AVR-PikF, AVR-Pia and the non-MAX effector AVR-Pii as previously described (De la Concepcion et al., 2022, 2018) (See materials and methods) and performed multicycle kinetics by flowing the effectors over a Biacore CM5 chip presenting amine-coupled RGA5, APB and Pikm-1 HMA domains (**Figure S6, Table S1**). As in previous reports, we observed strong binding of AVR-PikD to the Pikm-1 HMA (equilibrium dissociation constant (*K*_D_) = ∼10 nM) (De la Concepcion et al., 2018) and very low binding to RGA5 HMA (Cesari et al., 2022; Zhang et al., 2022) (**Figure 4 D, Figure S6**); neither the Pikm-1 HMA or RGA5 demonstrated any strong binding to AVR-PikC or AVR-PikF, as characterised by their rapid dissociation from the HMA (**Figure S6 A, Table S1**). Due to the weak binding of AVR-Pia to the HMAs relative to the Pikm-1/AVR-PikD interaction, higher concentrations (up to 50 µM) of AVR-Pia were flowed over the chip, which allowed us to measure the affinity of AVR-Pia for the RGA5 and APB HMAs at 26.8 and 32.9 µM, respectively (**Figure S6**). These values are in agreement with previous studies that have reported micromolar affinities between the RGA5 HMA and AVR1-CO39 and AVR-Pia effectors using isothermal titration calorimetry (ITC) or microscale thermophoresis (MST) (Guo et al., 2018; Liu et al., 2021; Ortiz et al., 2017; Zhang et al., 2022). Interestingly, the affinity of the interaction between RGA5 HMA and AVR-Pia is similar to the affinities observed for the interaction between RGA5 HMA and AVR-PikD, or Pikm-1 HMA and AVR-PikC/AVR-PikF, which do not to result in resistance in planta. A key similarity between these weaker HMA/effector interactions is the rapid dissociation rate of the effector from the HMA, indicative that the RGA5 HMA alone can facilitate, but not maintain, binding of the effector in vitro. This observation is particularly evident when compared to the binding of Pikm-1 to AVR-PikD in which the dissociation rate is considerably slower (**Figure S6**).

We observed high binding affinity of AVR-PikD, AVR-PikC and AVR-PikF for the APB mutant (**Figure 4 D**). As the effector does not dissociate appreciably from the HMA over the time of the experiment, we were unable to accurately calculate binding using multicycle kinetics (**Figure S7**). Therefore, to quantify the affinity between the APB and AVR-Pik effectors, we used single-cycle kinetics with a long dissociation phase, which allowed us to calculate a *K*_D_ of 0.31 nM, 2.95 nM, and 16.50 nM for AVR-PikD, AVR-PikC, and AVR-PikF respectively (**Figure 4 D, Table S1**).

The engineered APB mutant can bind AVR-Pik effectors with nanomolar affinity, and this strong binding correlates with the effector association and cell death response observed for Pikm-1^APB^ in planta. By contrast, AVR-Pia rapidly dissociates from all HMA domains, and this is correlated with weak/no response with Pikm-1^APB^ and Pikm-1^RGA5^ chimeras in planta. Taken together, these data suggest binding affinity to the HMA domain is key to recognition in the Pik system, with high affinity interfaces being essential for initiating a cell death response.

### The structural basis for interaction between the RGA5-APB HMA mutant and AVR-Pik

The Pikm-1 HMA and RGA5 HMA domains are essential for recognition of MAX effectors in their respective NLRs, however they have spatially distinct effector binding interfaces (De la Concepcion et al., 2018; Guo et al., 2018; Ortiz et al., 2017).

As the effector recognition interfaces of RGA5 and Pikm-1 HMA domains are different, we determined a crystal structure of the complex between the APB mutant and AVR-PikF, to validate our structural modelling of the RGA5 HMA and confirm we had engineered an AVR-Pik/Pik-HMA-like interface into RGA5 HMA. Using analytical gel filtration, we observed a peak shift after incubating purified AVR-PikF and APB proteins, indicative of stable complex formation (**Figure 5 A**). Following this, we used a co-expression approach to purify an APB/AVR-PikF complex, which was used to obtain crystals via sparse matrix screening (**Figure S8**). X-ray diffraction data were collected at the Diamond Light Source, Oxford, resulting in a 1.3Å dataset (**Table S2**) **(See details of crystallization and structure solution in materials and methods)**. The APB/AVR-PikF complex shares the same interface as Pikm-1 HMA/AVR-PikD and OsHIPP19/AVR-PikF complexes. Structural alignment of these complexes results in an R.M.S.D. of 0.51 Å and 0.39 Å, respectively (**Figure 5 B, D; Figure S9**). As predicted, each of the six mutations in the RGA5 HMA generated to facilitate AVR-Pik binding are located at the effector interface (**Figure 5 B; Figure S10**). These mutations are sufficient to generate an AVR-Pik binding interface in the RGA5 HMA distinct from that observed for AVR-Pia/AVR1-CO39 (**Figure 5 C**).

**Figure 5.**
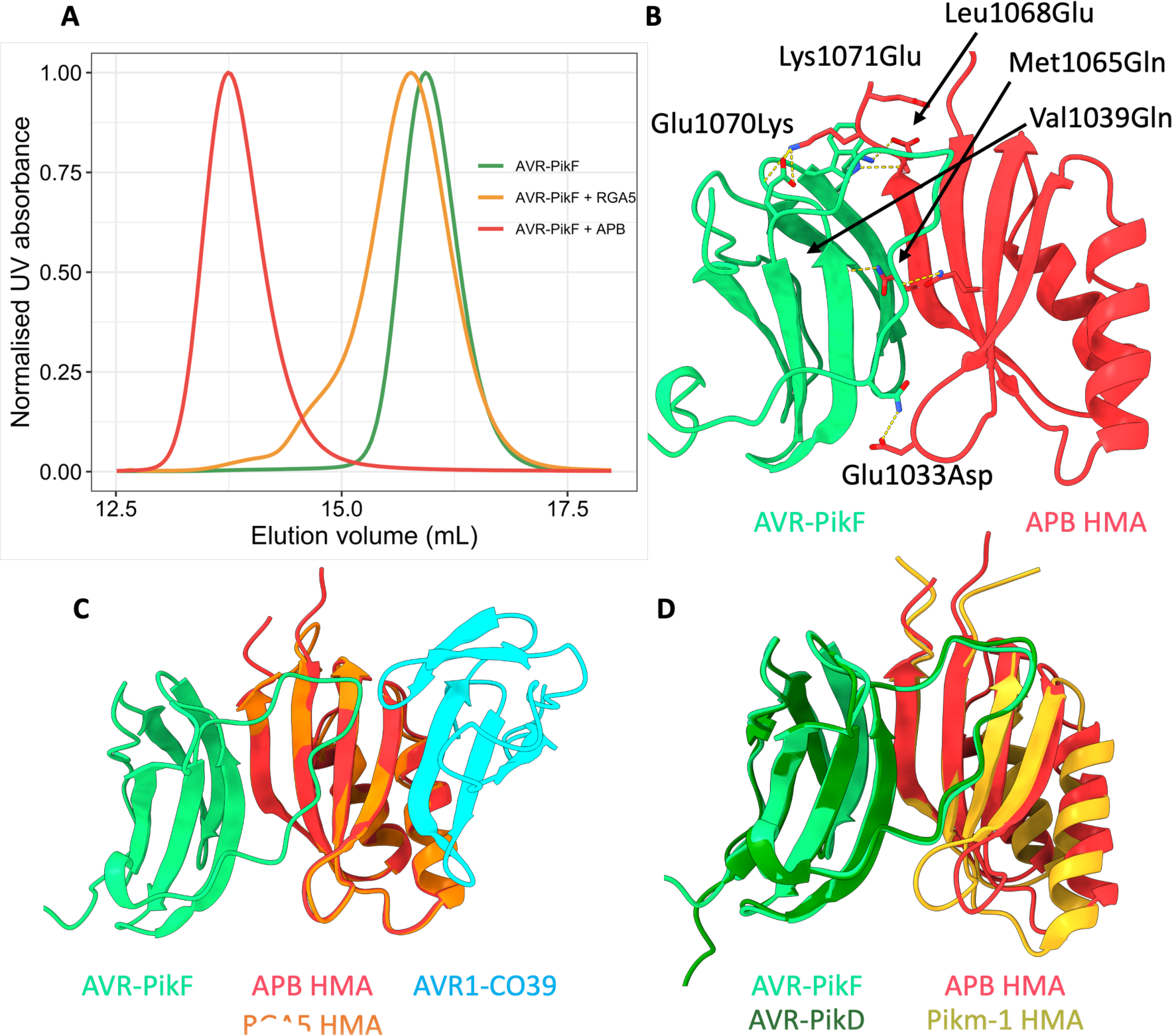
Six mutations in the RGA5 HMA reconstitutes a high affinity AVR-Pik binding interface akin to that of the Pik-1 HMA. **A)** Analytical size-exclusion chromatography of AVR-PikF with the RGA5 and APB HMA proteins. A mixture of AVR-PikF and APB HMA (red) elutes earlier than a mixture of RGA5 and AVR-PikF (orange) or AVR-PikF alone (green), indicative of complex formation between AVR-PikF and the APB HMA. **B)** The crystal structure of AVR-PikF in complex with the RGA5 APB HMA mutant (PDB: 8B2R). Mutations in RGA5, guided by the structure of the OsHIPP19/AVR-PikF complex, are shown forming contacts with AVR-PikF and are labelled. **C)** Superimposition of the crystal structures of the APB/AVR-PikF complex with the RGA5/AVR1-CO39 complex (PDB ID: 5ZNG) showing the swapped effector-binding interface of the APB HMA compared to the RGA5 HMA. **D)** Superimposition of the APB/AVR-PikF complex with the crystal structure of AVR-PikD bound to the Pikm-1 HMA domain (PDB ID: 6G10), showing the shared effector binding interface in these complexes.

## Discussion

Constitutive immune activation by the combination of incompatible NLRs through breeding or genetic engineering (hybrid necrosis) presents a bottleneck in plant breeding and evolution (Calvo-Baltanás et al., 2021; Chae et al., 2014; Ordon et al., 2021; Tran et al., 2017). Likewise, autoactivity due to engineering presents a bottleneck to strategies for NLR-mediated pathogen resistance (Kourelis et al., 2021; Maidment et al., 2021; Tamborski et al., 2022). The work presented here highlights the importance of factors outside of enhancing effector binding, such as considering the context of NLRs that act in pairs or networks, for the generation of new recognition specificities and NLR combinations without penalties imposed by constitutive immune activation.

### The helper allele Pikp-2 can accommodate for changes in the integrated domain of Pik-1 without triggering effector-independent cell death

We previously reported on the incompatibility of the Pikp and Pikm alleles (De la Concepcion et al., 2021b), highlighting the functional diversification of the Pik receptor pair and linking the specialisation of the Pik-2 receptor for its cognate sensor to an Asp230Glu polymorphism in the NB domain. Here we demonstrated the role of the HMA domain in compatibility of the Pik-1 sensor with Pik-2 the helper. When we introduced the HMA domain from Pikp-1 into Pikm-1 we observed autoactivity with Pikm-2 but not Pikp-2. Pikp-2^D230E^ and Pikm-2^E230D^ mutants, which flip the specialisation of each helper, swapped the compatibility of Pik-2 for Pik-1 mutants/chimeras.

Studies involving the swap of the Pikp integrated HMA for a non-co-evolved ancestral version (Białas et al., 2021) or the equivalent HMA domain of OsHIPP19 (Maidment et al., 2022) also showed that this caused autoimmunity, which was removed by mutation of the HMA outside of the effector binding interface, further supporting a mechanism for co-adaptation of the integrated HMA domain with other domains in the sensor Pik-1 and the helper Pik-2. This co-adaptation may have led to different thresholds of the helpers for the sensors. As such, we observe Pikp-2 to be more permissive of changes in the Pik-1 compared to Pikm-2.

Specific Pik pair combinations are more tolerant to changes in the integrated domain, facilitating engineering of expanded recognition that would otherwise result in constitutive cell death. By considering the context of the engineered receptor domain within the NLR pair, we present a novel approach to circumventing autoactive immune responses that can limit the potential of NLR engineering for novel disease resistance.

### Allelic mismatching provides new avenues for engineering disease resistance

The mismatching strategy reported here opens exciting avenues for the incorporation of new effector recognition motifs into the Pik system, and perhaps other paired NLR systems. Combining the Pikm-1 sensor with the Pikp-2 helper yielded a compatible receptor pair with greater ability to accept HMA modifications than the natural pairing of the Pikp-1/Pikp-2 or Pikm-1/Pikm-2 alleles. Mismatching of the Pik sensor/helper alleles allowed incorporation of the RGA5 HMA into the Pikm-1 backbone, without autoactivity. Notably, this strategy could be useful in areas such as the incorporation of VHH-nanobody fusions into Pikm-1 to allow for tailor-made NLRs (Kourelis et al., 2021). Indeed, in this study several Pikm-1/VHH-nanobody chimeras triggered constitutive cell death responses, indicative of autoactivity. Mismatching of the Pik alleles in conjunction with nanobody integration could allow for a greater number of successful integrations, streamlining this engineering strategy.

Recently, there have been several reports of engineering expanded effector recognition in integrated HMA domain containing NLRs (Cesari et al., 2022; De la Concepcion et al., 2019; Liu et al., 2021; Maidment et al., 2022; Zhang et al., 2022). In the RGA5/RGA4 system, recognition of AVR-Pib and AVR-PikD has been engineered into the RGA5 HMA domain, however this results in compromised AVR-Pia recognition (Cesari et al., 2022; Liu et al., 2021; Zhang et al., 2022). Furthermore, implementation of model-driven engineering of RGA5 into crop systems is challenging and has had variable success, with RGA5 mutants that exhibit expanded recognition in a *N. benthamiana* model not always translating to disease resistance in transgenic rice lines (Cesari et al., 2022; Zhang et al., 2022). In parallel, engineering of the Pikp-1 HMA to respond to previously unrecognised AVR-Pik variants has been shown in *N. benthamiana* assays and transgenic rice lines (Maidment et al., 2022). Interestingly, full replacement of the integrated HMA for the HMA domain of OsHIPP19 caused autoimmunity, which was removed by mutation of the HMA. However, this is not always possible as the approach benefitted from knowledge of the NLR-ID/effector complex (Białas et al., 2021; De la Concepcion et al., 2019; Maidment et al., 2022). Whether engineering facilitated by allelic mismatching of the Pik pair can provide resistance in transgenic rice lines is yet to be tested and is an important next step to demonstrate the use of this approach for translating recognition in plant models to resistance in crops.

### The Pik system relies on high affinity effector binding to activate defence responses in planta

We demonstrate the RGA5 HMA domain can be integrated into the Pikm-1 backbone and engineered to recognise AVR-Pik, including variants not recognised by wildtype Pikm-1. As shown by our biophysical and structural analysis, the six mutations introduced in the APB mutant of RGA5 HMA domain, based on the host target OsHIPP19, recapitulate a functional AVR-Pik binding recognition interface. These data highlight the power of host target-guided design of NLR-ID baits for engineering recognition.

While low levels of cell death were observed, neither the Pikm-1^RGA5^ nor Pikm-1^APB^ responded to AVR-Pia at a level comparable with RGA5/RGA4. It is possible the AVR-Pia/AVR1-CO39 interface is occluded in the Pikm-1^RGA5^ chimera, and co-IP with the APB mutant did not show association in planta. However, we speculate the lack of AVR-Pia recognition *N. benthamiana* may more likely be due to a lower affinity of the effector for the HMA domain, as we were able to observe some weak cell death when Pikm-1^RGA5^ or Pikm-1^APB^ co-expressed with Pikp-2 was challenged with AVR-Pia. Previous studies have benchmarked the affinity of AVR-Pia/AVR1-CO39 for the RGA5 HMA domain in the micromolar range (Guo et al., 2018; Ortiz et al., 2017), while AVR-Pik effectors bind their cognate integrated HMA domains with nanomolar affinity (for interactions that result in cell death responses). It remains unclear why the binding affinities of the Pik HMA and RGA5 HMA domains for their cognate effectors differ so significantly. However, the RGA5 post-LRR region, which contains the HMA domain, also contains several other small uncharacterised domains (**Figure S11**), which may contribute to effector binding. Indeed, AVR-Pia is known to associate with regions of the RGA5 receptor outside of the HMA domain (Ortiz et al., 2017). However, RGA5 receptors where the HMA domain has been deleted are unable to respond to AVR-Pia in cell death assays (Cesari et al., 2013). Furthermore, a recent studies engineering AVR-Pib and AVR-Pik recognition in RGA5, respectively, showed mutations outside of the HMA influenced effector recognition in planta (Liu et al., 2021; Zhang et al., 2022). Collectively, available data suggest that RGA5-mediated effector recognition requires the HMA domain, but alone it is not sufficient for effector recognition, and works together with other regions at the RGA5 C-terminus. Inclusion of these additional regions from RGA5 into the Pik receptor backbone alongside the RGA5 HMA domain could support AVR-Pia recognition, but equally so, may affect Pik sensor/helper compatibility. Certainly, the additional complexity of the RGA4/RGA5 system makes engineering this receptor pair more challenging compared to the Pik NLRs.

Modification of plant NLRs has proven challenging due to the lack of understanding of the context of NLRs as part of complex systems. In this study, we demonstrate a new avenue for NLR-mediated resistance engineering that exploits the allelic diversity in the Pik NLR pair to allow for generation of receptors with expanded recognition specificities that would otherwise result in constitutive cell death. Our structural, biophysical and in planta analyses demonstrate the Pik system requires a high affinity effector binding interface to allow for binding to translate to defence, and as a single domain, the RGA5 HMA domain appears to lack the affinity for AVR-Pia to facilitate a robust Pik chassis-mediated cell death response. However, our engineering of RGA5 HMA to recognise AVR-Pik from within the Pikm-1 chassis highlights the strengths of this system for engineering; only a single high-affinity interface needs to be present to mediate effector recognition, making the Pik system a simple but efficient means for generating bespoke NLR resistance. This work lays the foundation for the incorporation of new effector recognition motifs into the Pik system and is a key advance towards the development of designer NLRs that can be tailored to specific secreted pathogen signatures.

## Acknowledgements

We thank the Diamond Light Source, UK (beamline i04 under proposal 25108) for access to X-ray data collection facilities and Dave Lawson from the JIC Protein Crystallography Platform for expert technical assistance during data collection. We thank Dan McLean for technical advice with the besthr R package. We further thank all members of the BLASTOFF team at the Sainsbury Laboratory, John Innes Centre and Iwate Biotechnology Research Center. We especially thank Vincent Were for providing PWL2 constructs. This work was supported by UKRI Biotechnology and Biological Sciences Research Council (BBSRC) Norwich Research Park Biosciences Doctoral Training Partnership, (grant BB/M011216/1); the UKRI BBSRC, UK (grants BB/P012574/1, BBS/E/J/000PR9797) the European Research Council (ERC proposal 743165), the ERAMUS+ training programme, the John Innes Foundation and the Gatsby Charitable Foundation.

---

**Supplementary figure 1.**
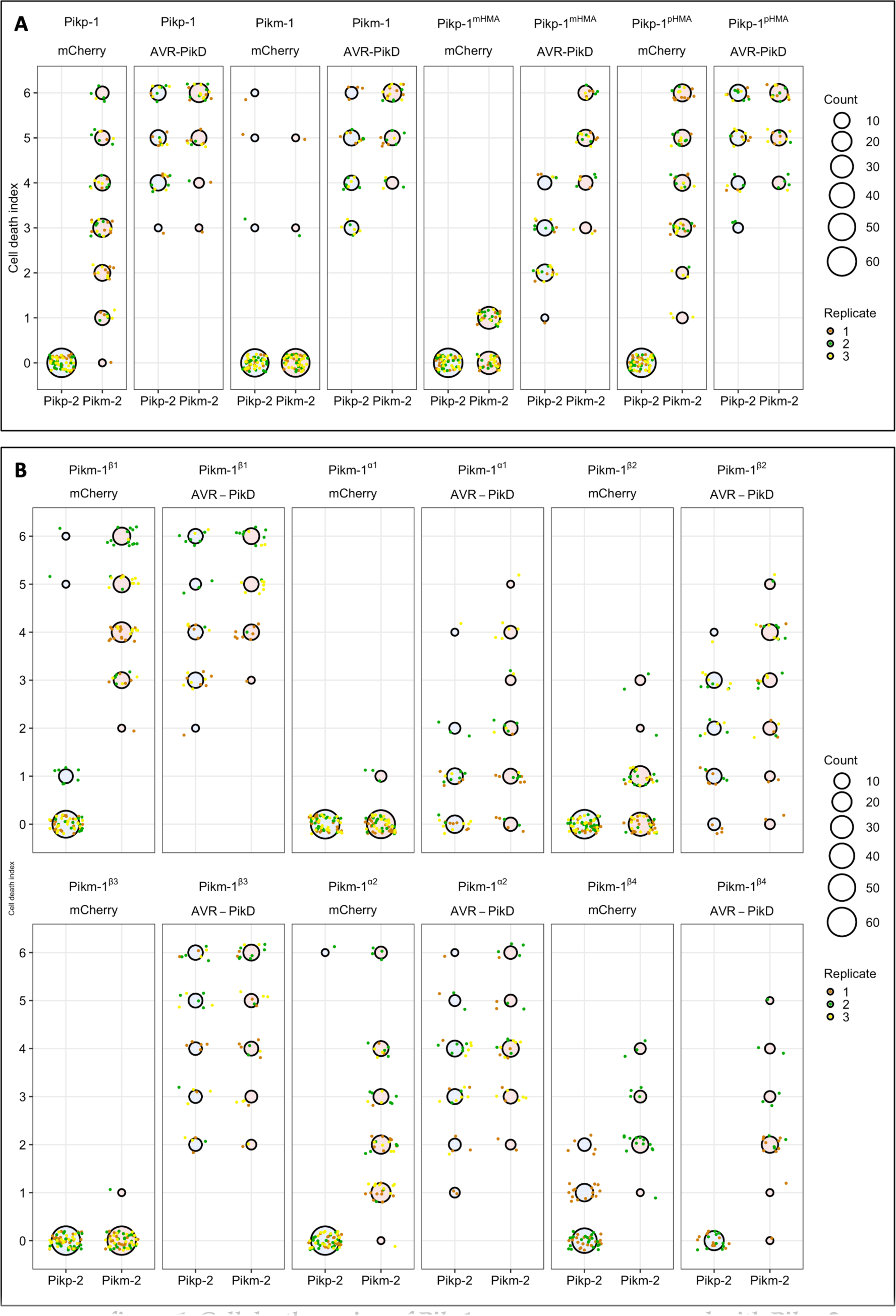
Cell death scoring of Pik-1 chimeras co-expressed with Pikp-2 and Pikm-2 in *N. benthamiana*. **A)** Scoring of Pikp-1^mHMA^ and Pikm-1^pHMA^ chimeras when co-expressed with Pikp-2 or Pikm-2 and either AVR-PikD or mCherry. **B)** Scoring of Pikm-1 chimeras carrying different secondary structures from the Pikp-1 HMA when co-expressed with Pikp-2 or Pikm-2 and either AVR-PikD or mCherry. Scoring is represented as dot plots. The total number of repeats was 60 per sample for both **A** and **B**. For each sample, all the data points are represented as dots with a distinct colour for each of the three biological replicates; these dots are jittered around the cell death score for visualisation purposes. The size of the central dot at each cell death value is proportional to the number of replicates of the sample with that score. Statistical analysis of these results is shown in **Appendix 1 B**.

**Supplementary figure 2.**
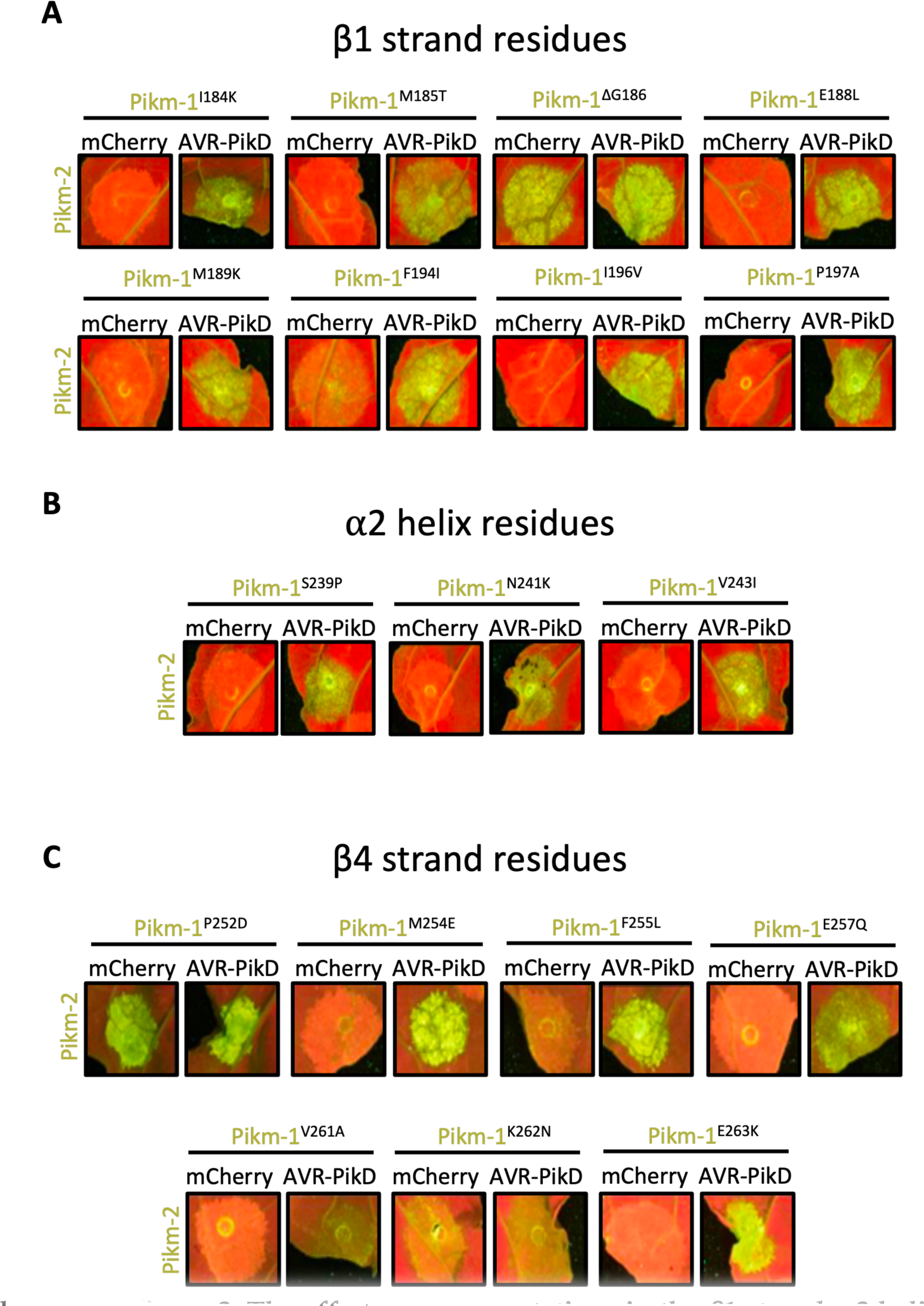
The effect of point mutations in the β1 strand, α2 helix, and β4 strand secondary structures of the Pikm-1 HMA domain, and their effect on compatibility with the Pikm-2 helper in *N.* *benthamiana*. Point mutations are substituted with the corresponding residue in the Pikp HMA. A) Individual point mutations of the residues in the β1 strand. B) Individual point mutations of the residues in the α2 helix. C) Individual point mutations of the residues in the β4 strand. Quantification and statistical analysis of these results are shown in **Figure S3**, Appendices 1 D, E, F.

**Supplementary figure 3.**
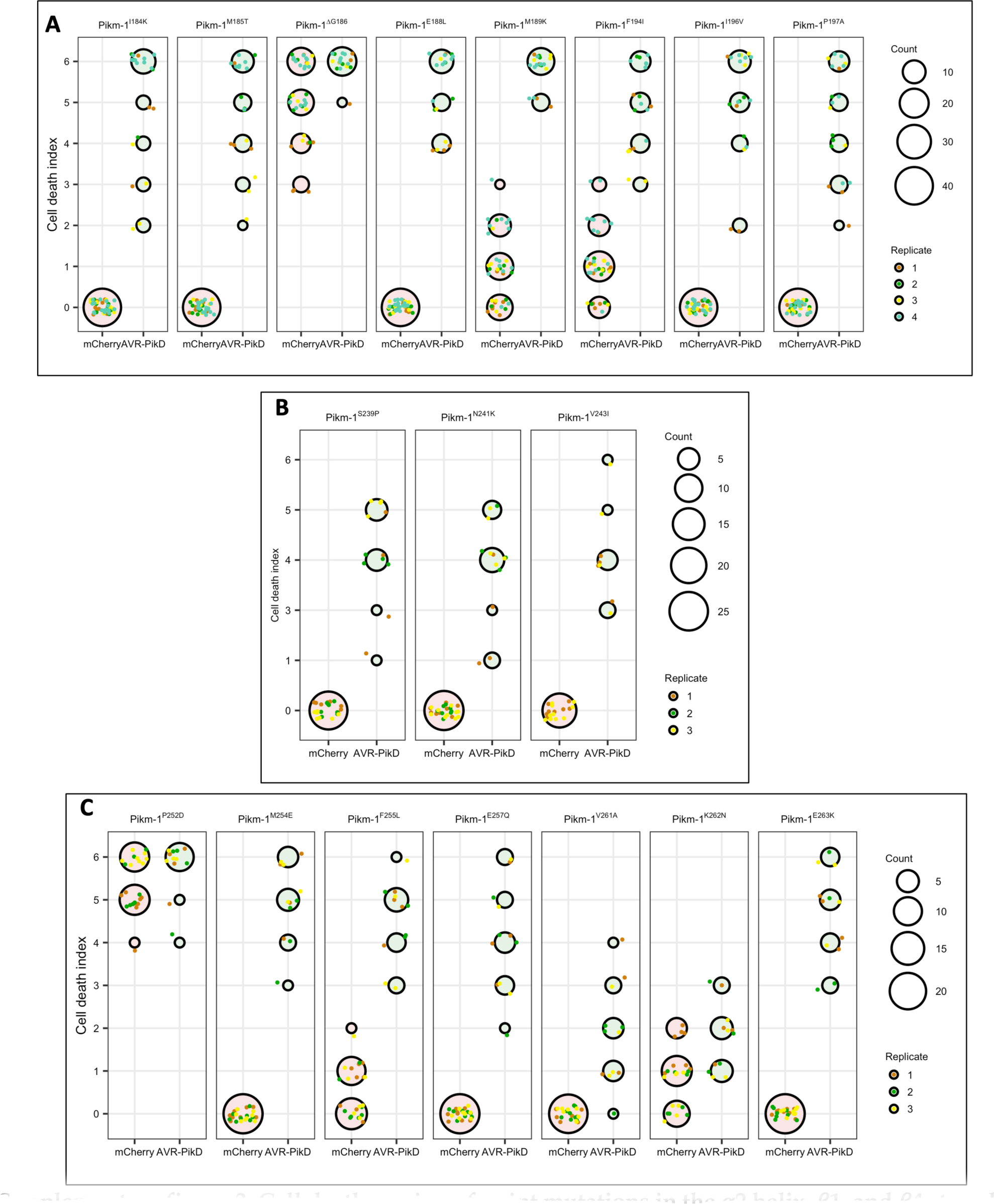
Cell death scoring of point mutations in the α2 helix, β1, and β4 strands of the Pikm-1 HMA domain when expressed with the Pikm-2 helper in *N. benthamiana*. **A)** Residues belonging to the β1 strand. B) Residues belonging to the α2 helix. C) Residues belonging to the β4 strand. Scoring is represented as dot plots. The total number of repeats was 40 per sample. For each sample, all the data points are represented as dots with a distinct colour for each of the three biological replicates; these dots are jittered around the cell death score for visualisation purposes. The size of the central dot at each cell death value is proportional to the number of replicates of the sample with that score. Quantification and statistical analysis of these results are shown in Appendices 1 D, E, F.

**Supplementary figure 4.**
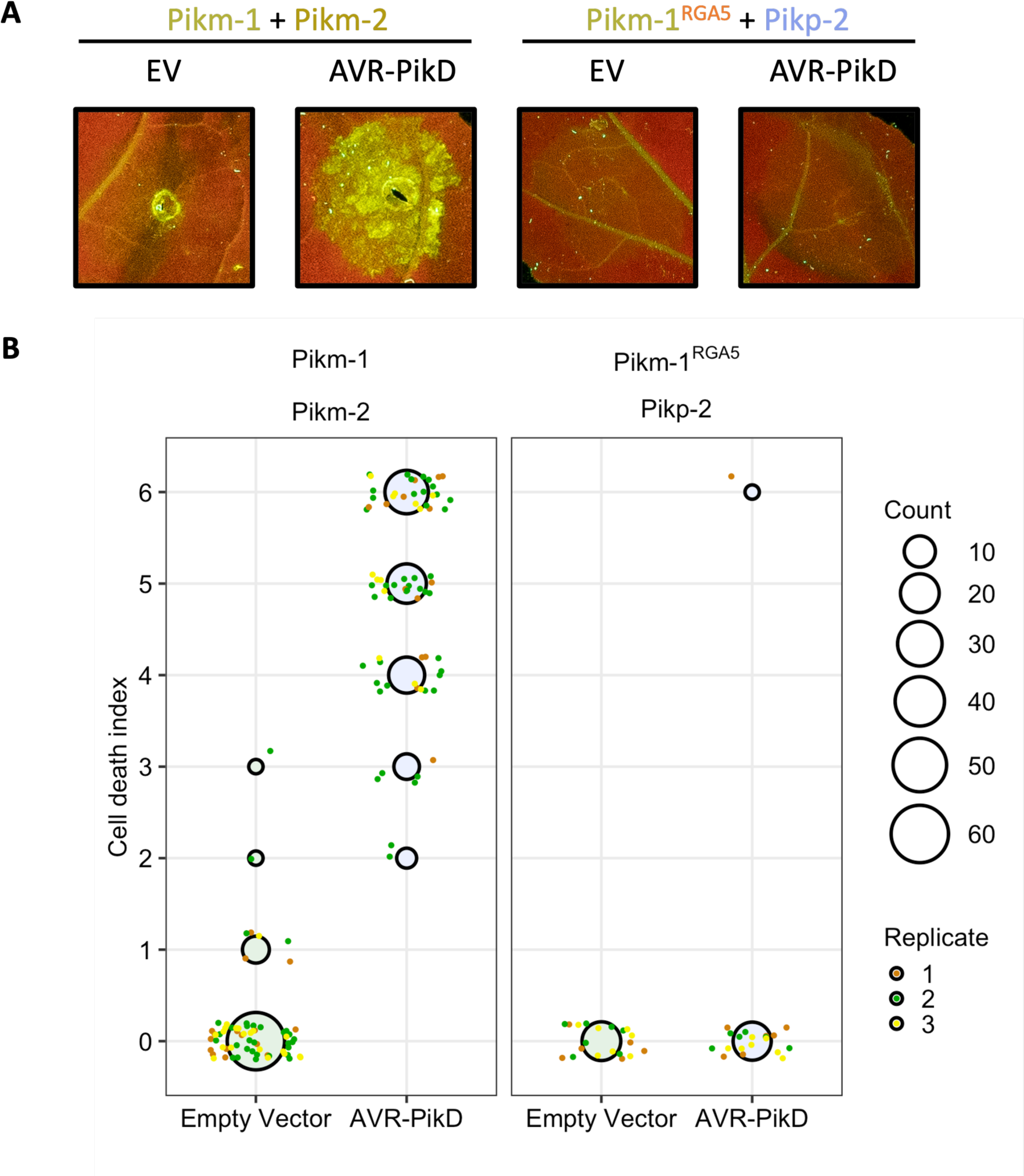
The Pikm-1^RGA5^ chimera does not respond to AVR-PikD in *N. benthamiana*. **A)** Co-expression of the Pikm-1^RGA5^ chimera with Pikp-2 and AVR-PikD in *N. benthamiana* leaves does not result in cell death. Wild-type Pikm-1 and Pikm-2 co-expressed with AVR-PikD shown as positive control **B)** Cell death scoring of **A)** represented as dot plots. For each sample, all the data points are represented as dots with a distinct colour for each of the three biological replicates; these dots are jittered around the cell death score for visualisation purposes. The size of the central dot at each cell death value is proportional to the number of replicates of the sample with that score. Statistical analyses of these results are shown in **Appendix 1 H**.

**Supplementary figure 5.**
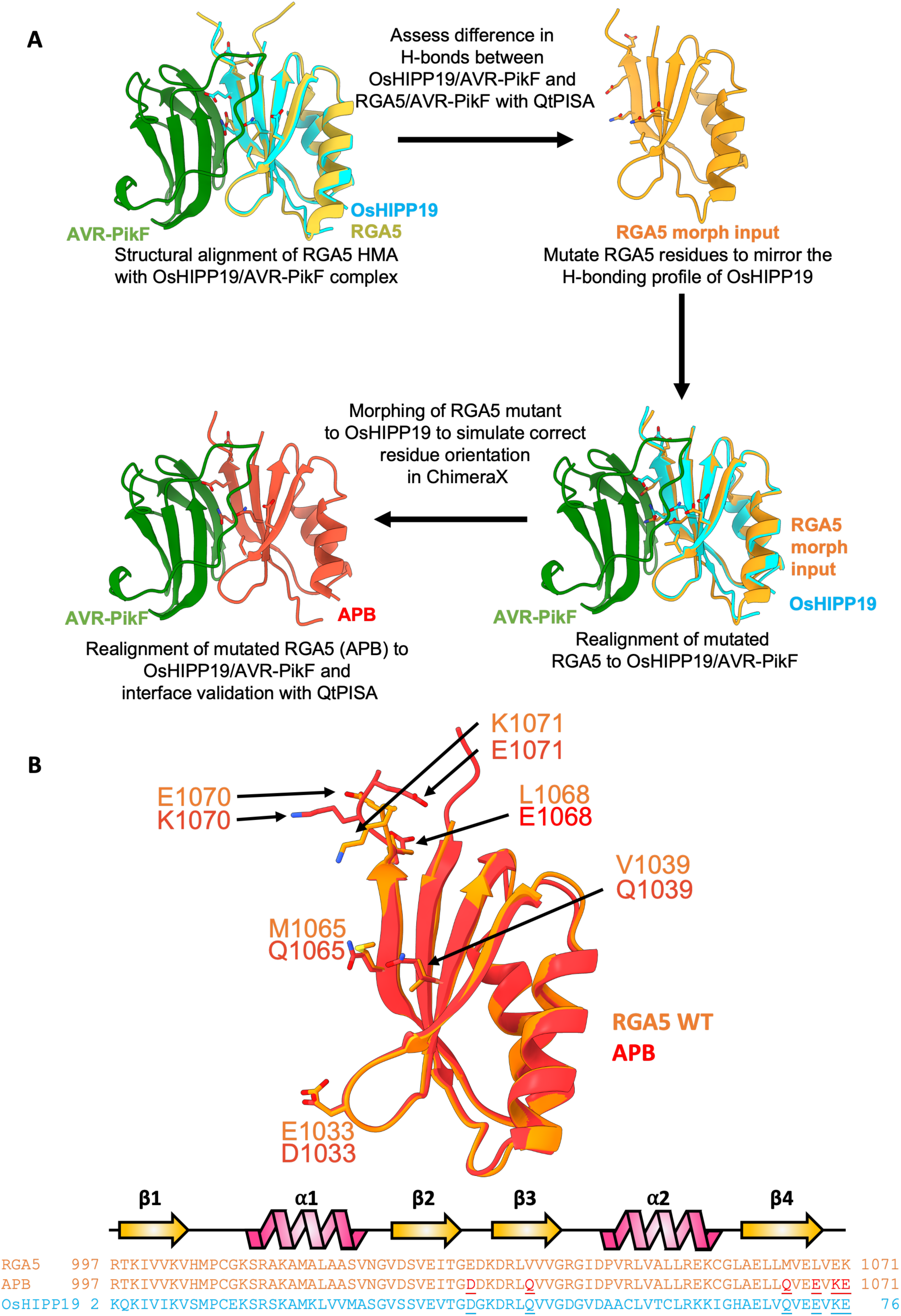
Structure-guided engineering of RGA5 using OsHIPP19 as a template to generate the APB mutant. **A)** Modelling pipeline using QtPISA and ChimeraX to make the RGA5 APB mutant using OsHIPP19 as a template. **B)** Structural alignment of the RGA5 HMA (PDB: 5ZNG) with the APB mutant with a sequence alignment highlighting the changes informed by OsHIPP19.

**Supplementary figure 6.**
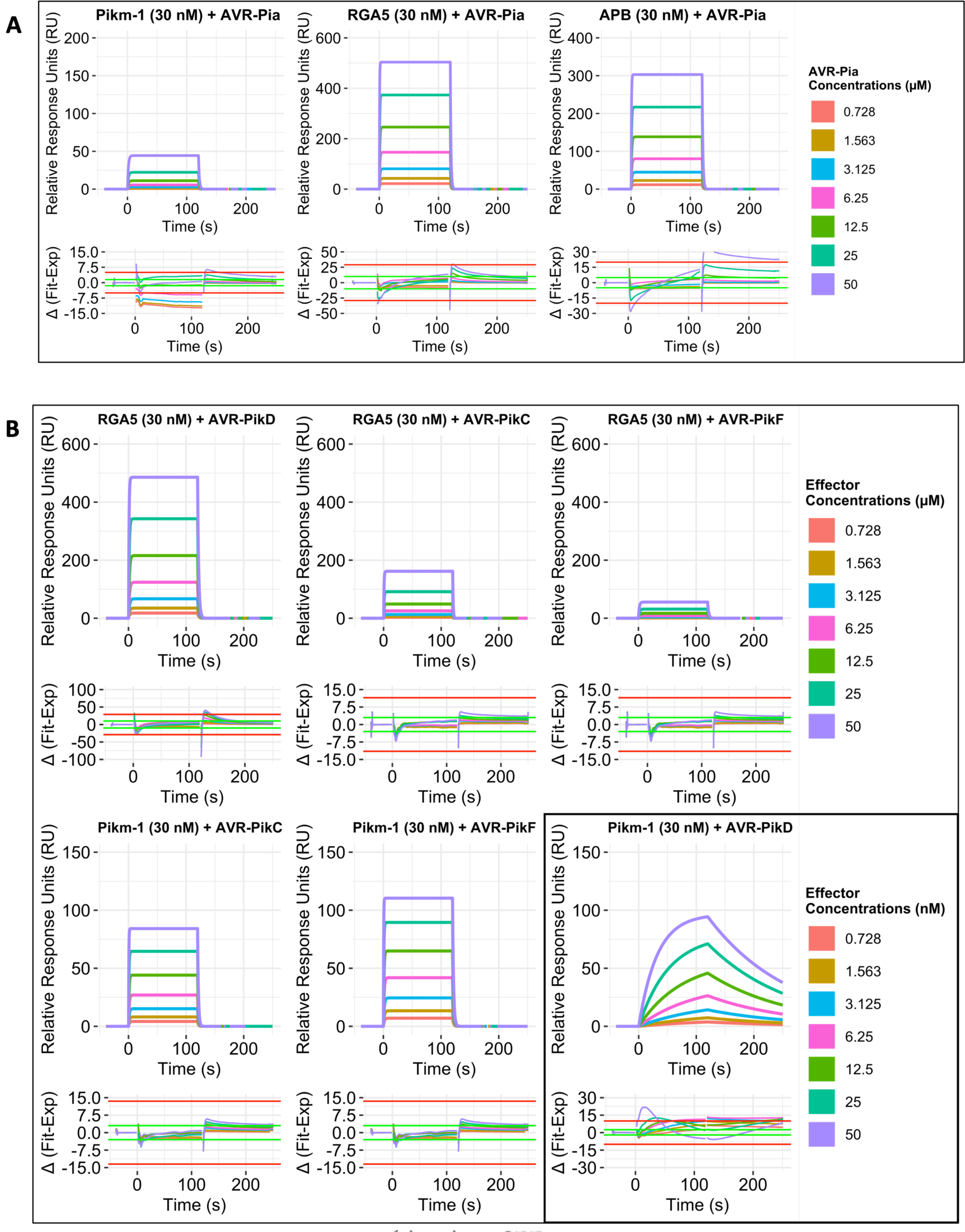
Multicycle kinetics SPR sensograms capturing effector/HMA interactions. Effectors were flowed over the HMA-bound CM5 chip at 8 concentrations (0 – 50 µM), with the exception of the strong Pikm-1 / AVR-PikD interaction where 0 – 50 nM of AVR-PikD was used. Kinetic and binding parameters were calculated using a 1:1 binding model. Residual graphs are shown under the sensograms, with data between the red lines being deemed reliable. **A)** Sensograms of the Pikm-1, RGA5 and APB HMA domains with AVR-Pia. **B)** Sensograms of RGA5 and Pikm-1 HMA domains with AVR-Pik effectors.

**Supplementary figure 7.**
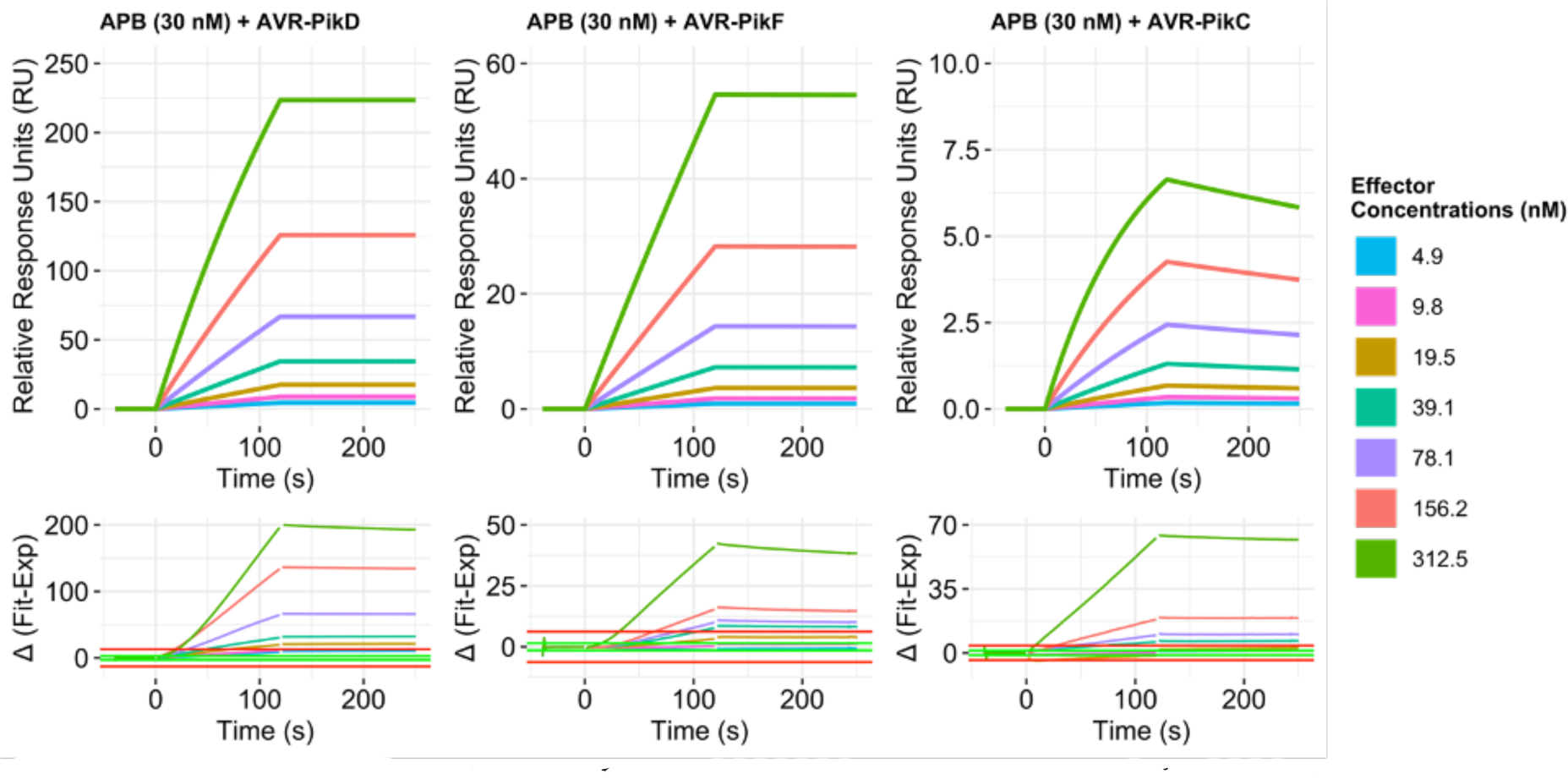
Multicycle kinetics SPR sensograms of the APB mutant with AVR-Pik variants. Effectors were flowed over the HMA-bound CM5 chip at 7 concentrations (4.9 – 312.5 nM). Kinetic and binding parameters were calculated using a 1:1 binding model. Residual graphs are shown under the sensograms, with data between the red lines being deemed reliable. Due to the high affinity of the effectors for the HMA, the effectors failed to dissociate from the APB chip between runs. This is reflected in the poor fit as described by the residuals, and the decreasing relative response as seen between the samples.

**Supplementary figure 8.**
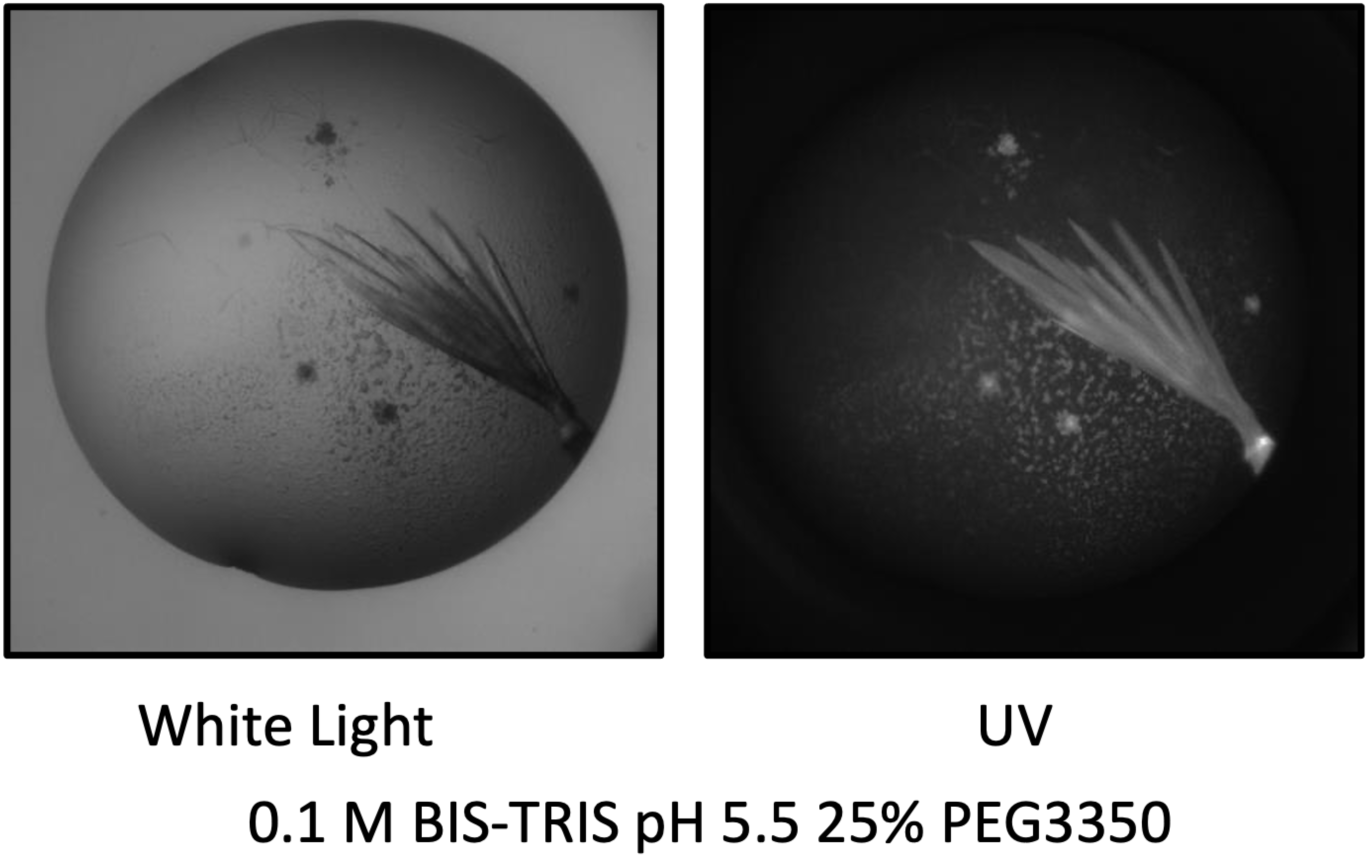
Crystallisation of the APB/AVR-PikF HMA complex. APB/AVR-PikF HMA complex crystals formed in ShotGun 1 sparse matrix screen (Molecular Dimensions) well B1 (0.1 M BIS-TRIS pH 5.5, 25% PEG3350) after 10 days.

**Supplementary figure 9.**
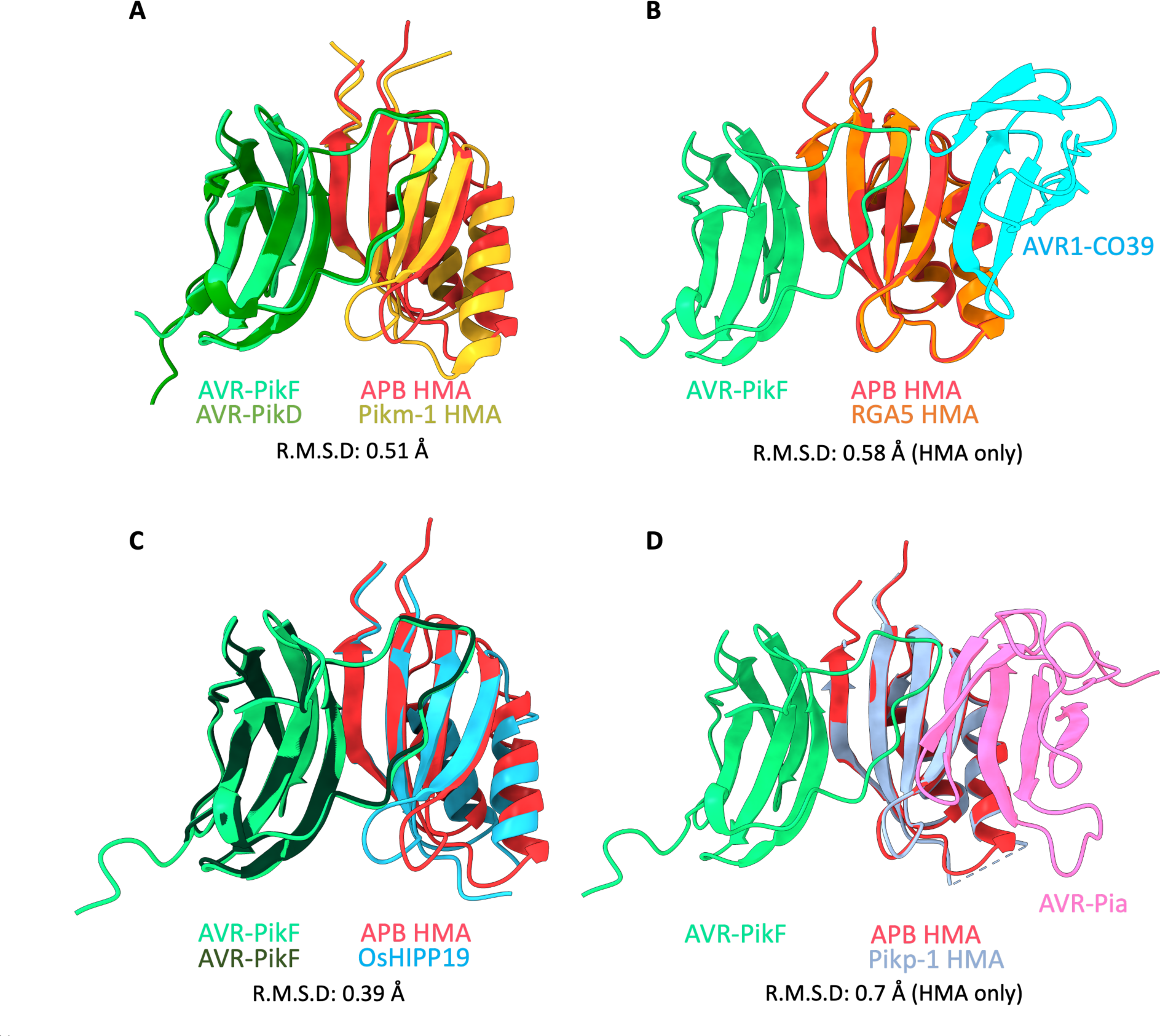
Superimposition of the crystal structure of APB HMA/AVR-PikF complex with other MAX effector/HMA complexes. **A)** Superimposition with Pikm-1 HMA/AVR-PikD (PDB ID: 6G10). **B)** Superimposition with RGA5 HMA/AVR1-CO39 (PDB ID: 5ZNG). **C)** Superimposition with OsHIPP19/AVR-PikF (PDB ID: 7B1I). **D)** Superimposition with Pikp-1 HMA/AVR-Pia (PDB ID: 6Q76).

**Supplementary figure 10.**
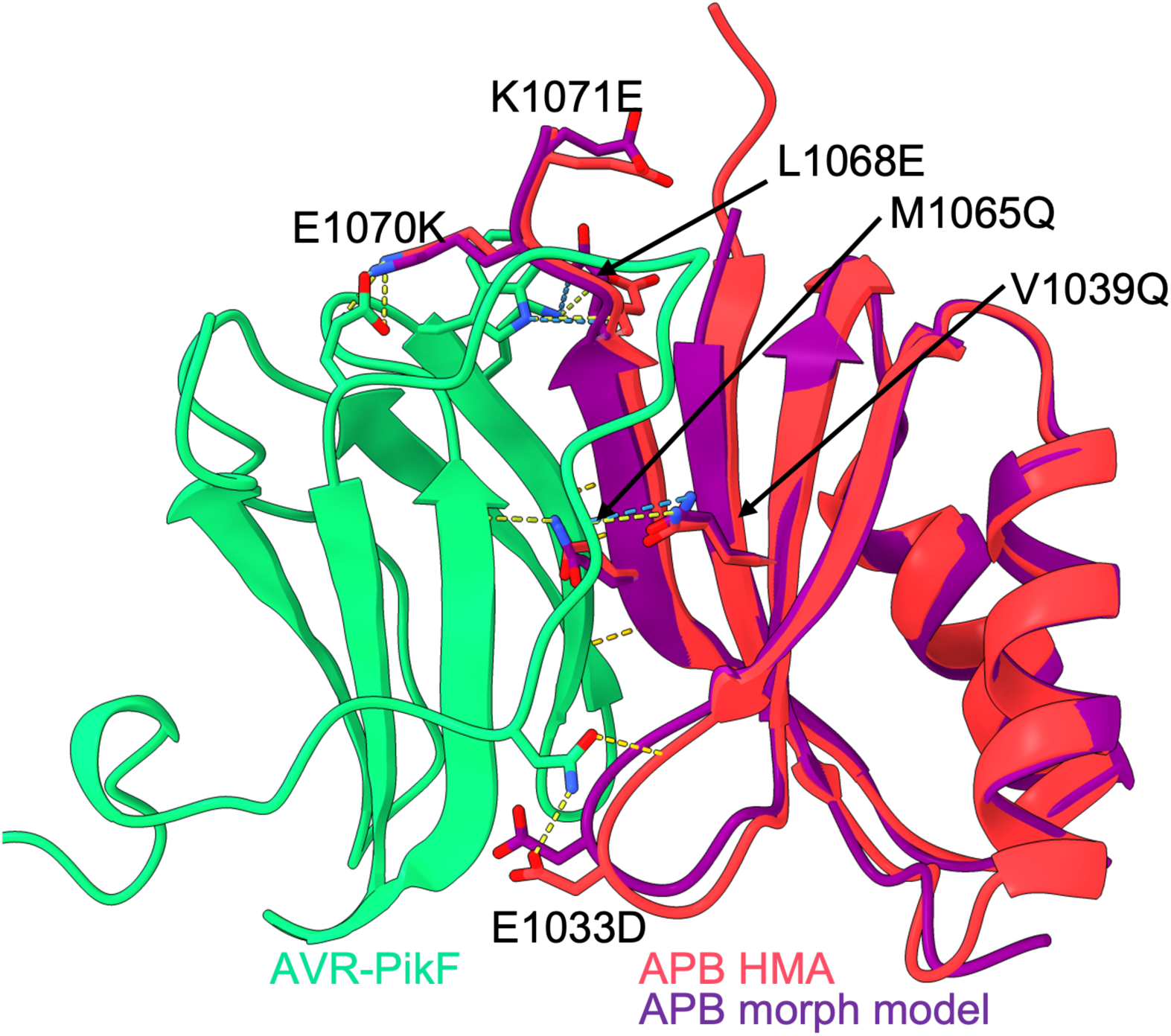
Superimposition of the crystal structure of APB HMA/AVR-PikF complex with the morph model generated from an OsHIPP19 template. Residues of RGA5 that were mutated based on OsHIPP19 to create the APB mutant are shown.

**Supplementary figure S11.**
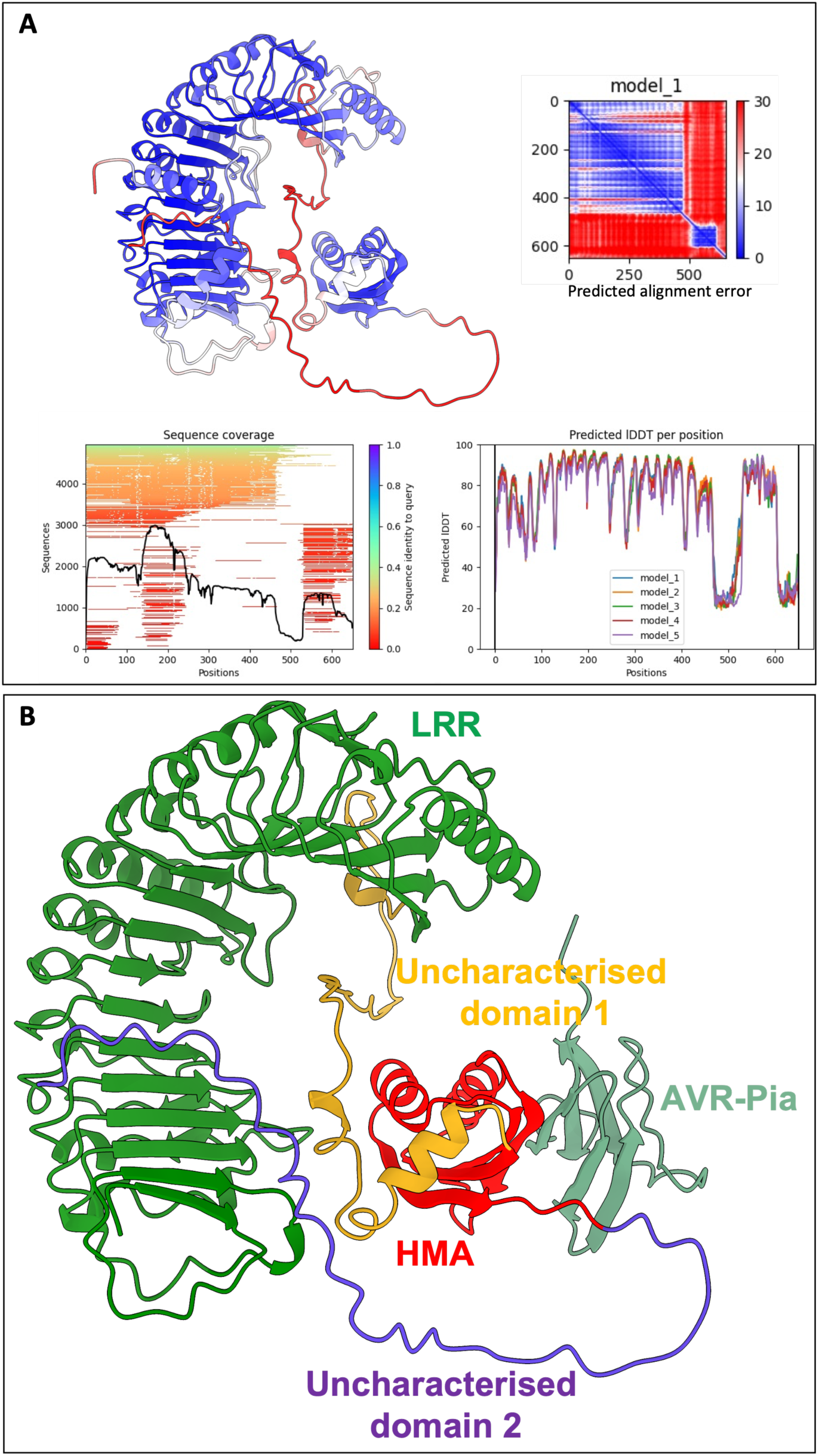
AlphaFold2 prediction of the C-terminal domains of RGA5 describes previously uncharacterised domains. **A)** AlphaFold2 v2.1 (Jumper et al., 2021) (as implemented in the AlphaFold ColabFold (Mirdita et al., 2022)) prediction of the RGA5 C-terminus. The RGA5 model is coloured by residue position confidences, with blue indicating high confidence and red low confidence. The LRR and HMA domains appear well predicted, however additional regions at the C-terminus lack confidence. **B)** Colouring of the RGA5 AlphaFold2 model to highlight the different domains present, with AVR-Pia (olive green) superimposed at the predicted HMA interface (from PDB ID: 6Q76).

**Supplementary table 1.**
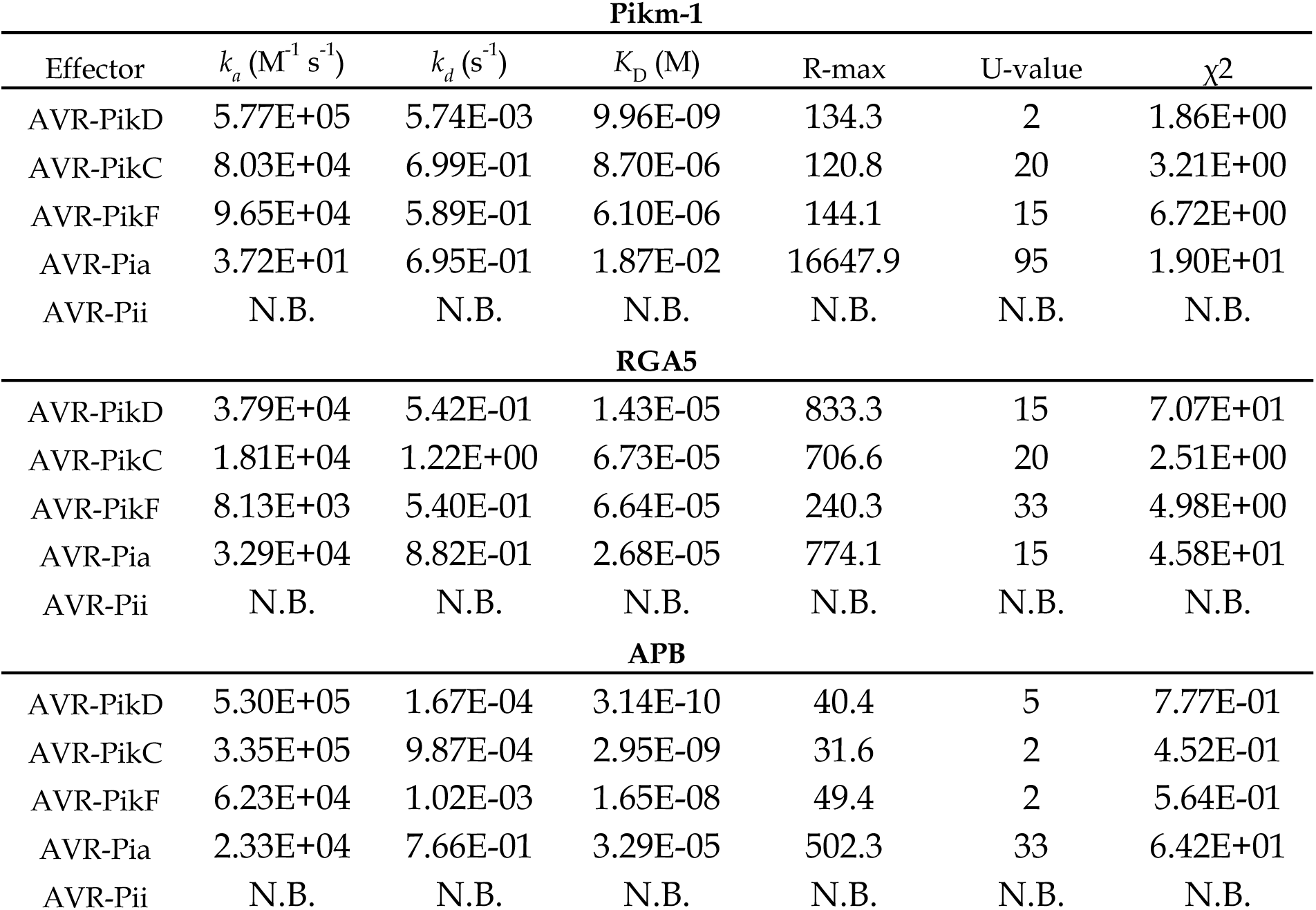
Full kinetic parameters for HMA-effector 557 interactions as measured by SPR.

**Supplementary table 2.** Data collection and refinement statistics APB/AVR-PikF

| APB/AVR-PikF |  |
| --- | --- |
| **Data collection statistics** |  |
| Wavelength (Å) | 0.979 |
| Space group | *P* 2<sub>1</sub> 2<sub>1</sub> 2<sub>1</sub> |
| Cell dimensions | 31.91 57.90 76.66 |
| *a*, *b*, *c* (Å) |  |
| Resolution (Å)* | 46.24 – 1.22 (1.24 – 1.22) |
| *R*<sub>merge</sub> (%) | 6.9 (188.4) |
| *I*/σ*I* | 14.5 (0.7) |
| Completeness (%) |  |
| Overall | 99.4 (92.0) |
| Anomalous | 99.0 (88.6) |
| Unique reflections | 42985 (1915) |
| Redundancy |  |
| Overall | 11.5 (5.2) |
| Anomalous | 6.0 (2.7) |
| CC(1/2) (%) | 100 (32.4) |
| **Refinement and model statistics** |  |
| Resolution (Å) | 46.24 – 1.22 (1.24 – 1.22) |
| *R*<sub>work</sub>/*R*<sub>free</sub> (%) | 14.9 / 20.2 |
| No. atoms | 1516 |
| Protein | 1324 |
| Ligand | 9 |
| Ion | 2 |
| Water | 181 |
| B-factors |  |
| Protein | 18.0 |
| Ligand | 33.6 |
| Ion | 22.1 |
| Water | 27.2 |
| R.m.s deviations |  |
| Bond lengths (Å) | 0.0129 |
| Bond angles (°) | 1.88 |
| Ramachandran plot (%)** |  |
| Favoured | 97.50 |
| Allowed | 1.88 |
| Outliers | 0.62 |
| MolProbity Score | 1.48 |

\*The highest resolution shell is shown in parenthesis.
\*\*As calculated by MolProbity

## Materials and Methods

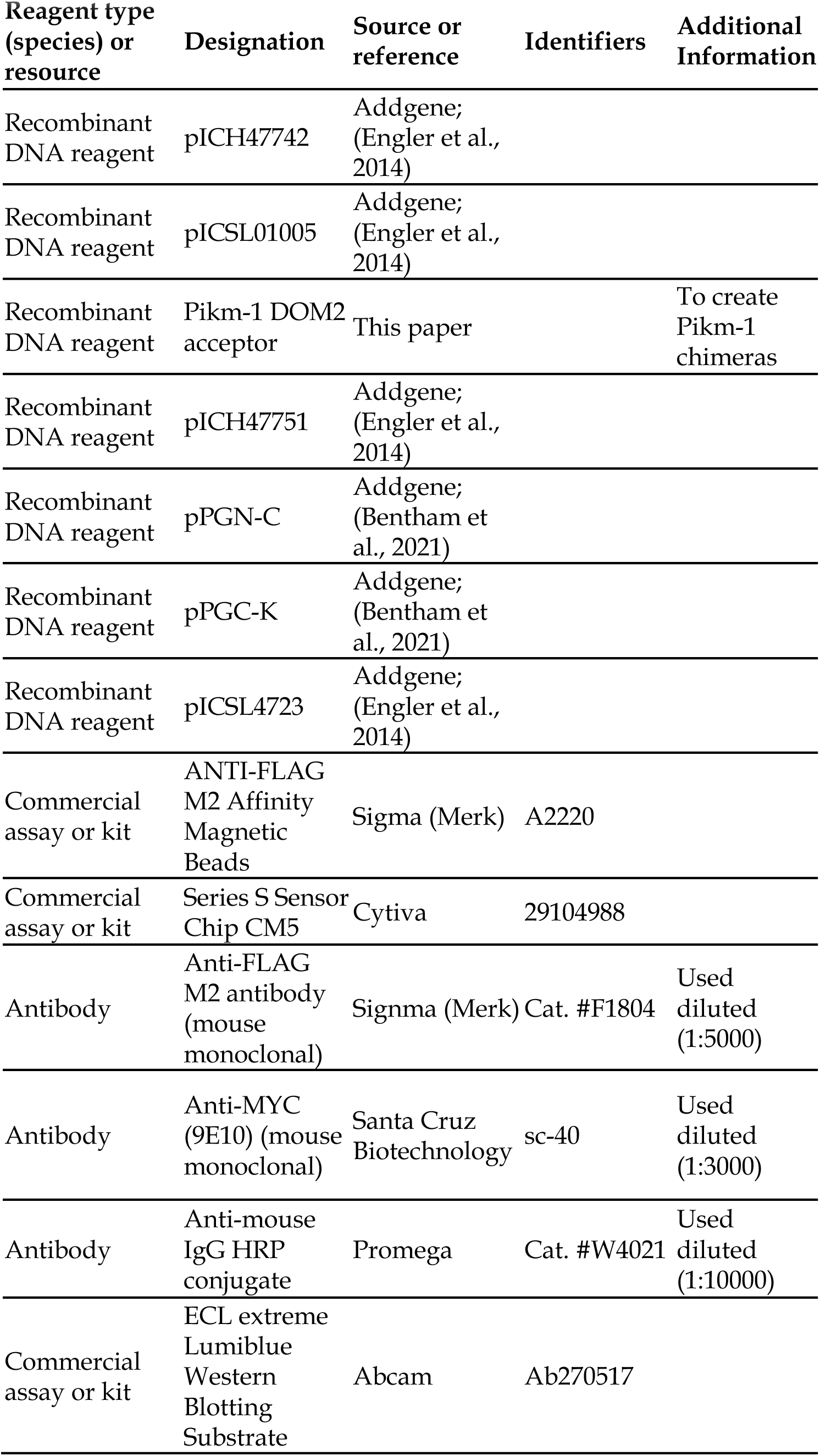
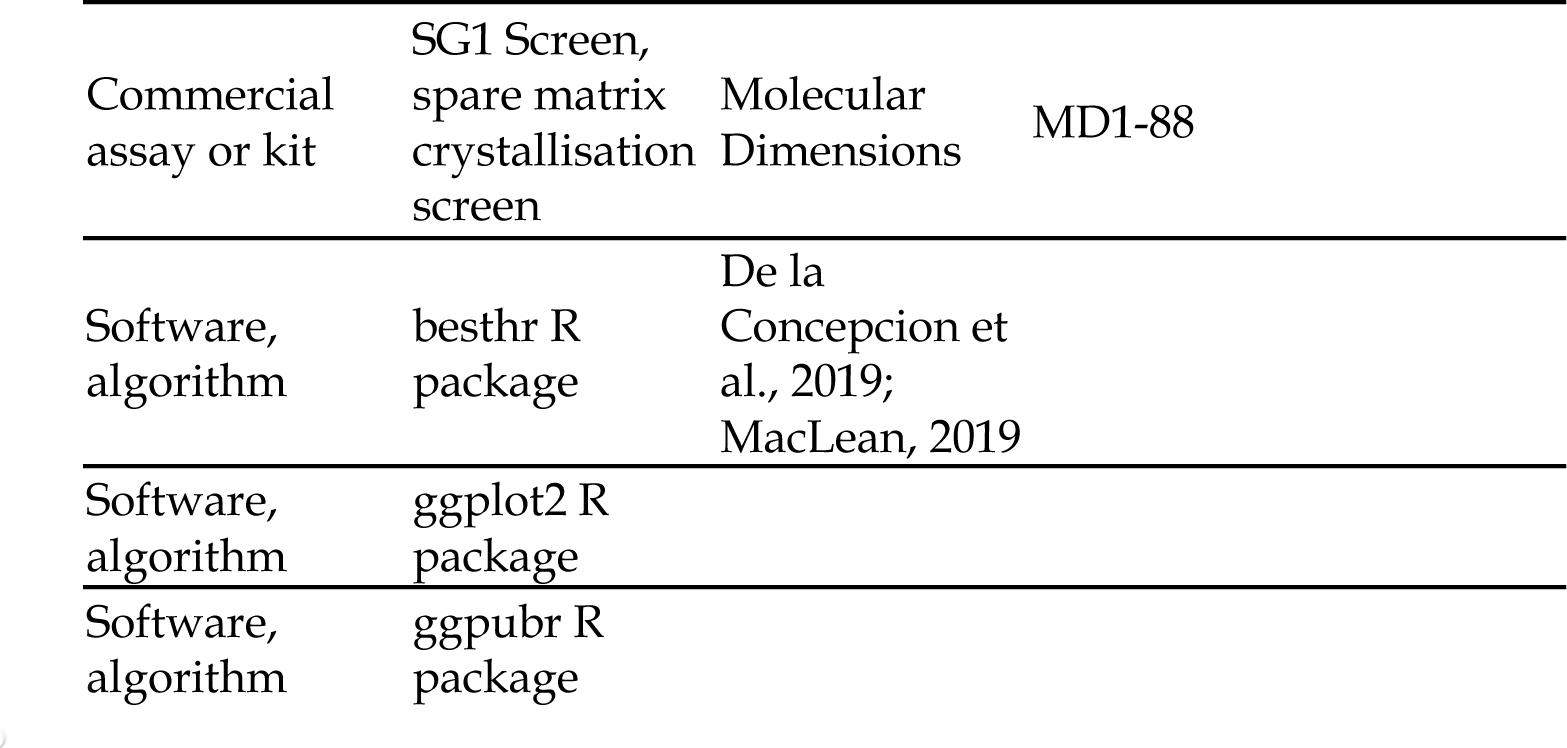
Key resources table

**Materials and Methods**565 **Key resources table**
| Reagent type (species) or resource | Designation | Source or reference | Identifiers | Additional Information |
| --- | --- | --- | --- | --- |
| Recombinant DNA reagent | pICH47742 | Addgene; (Engler et al., 2014) |  |  |
| Recombinant DNA reagent | pICSL01005 | Addgene; (Engler et al., 2014) |  |  |
| Recombinant DNA reagent | Pikm-1 DOM2 acceptor | This paper |  | To create Pikm-1 chimeras |
| Recombinant DNA reagent | pICH47751 | Addgene; (Engler et al., 2014) |  |  |
| Recombinant DNA reagent | pPGN-C | Addgene; (Bentham et al., 2021) |  |  |
| Recombinant DNA reagent | pPGC-K | Addgene; (Bentham et al., 2021) |  |  |
| Recombinant DNA reagent | pICSL4723 | Addgene; (Engler et al., 2014) |  |  |
| Commercial assay or kit | ANTI-FLAG M2 Affinity Magnetic Beads | Sigma (Merk) | A2220 |  |
| Commercial assay or kit | Series S Sensor Chip CM5 | Cytiva | 29104988 |  |
| Antibody | Anti-FLAG M2 antibody (mouse monoclonal) | Signma (Merk) | Cat. #F1804 | Used diluted (1:5000) |
| Antibody | Anti-MYC (9E10) (mouse monoclonal) | Santa Cruz Biotechnology | sc-40 | Used diluted (1:3000) |
| Antibody | Anti-mouse IgG HRP conjugate | Promega | Cat. #W4021 | Used diluted (1:10000) |
| Commercial assay or kit | ECL extreme Lumiblu Western Blotting Substrate | Abcam | Ab270517 |  |

|  |  |  |  |
| --- | --- | --- | --- |
| Commercial assay or kit | SG1 Screen, spare matrix crystallisation screen | Molecular Dimensions | MD1-88 |
| Software, algorithm | besthr R package | De la Concepcion et al., 2019; MacLean, 2019 |  |
| Software, algorithm | ggplot2 R package |  |  |
| Software, algorithm | ggpubr R package |  |  |

### Gene cloning – in planta expression

For expression in planta, full length Pikp-1 and Pikm-1, and relevant mutants, were cloned with a 6xHIS/3xFLAG tag into the pICH47742 plasmid, full length Pikp-2 and Pikm-2 were cloned into pICH47751 with a C-terminal 6xHA tag and Pikp-2^D230E^ and Pikm-2^E230D^ mutants were generated by site-directed mutagenesis as previously described (De la Concepcion et al., 2021b, 2018).

### Gene cloning – Generation of in planta expression constructs

To generate the Pikm-1 DOM2 acceptor, the Pikm sequence was domesticated of *BsaI* and *BbsI* restriction sites to allow compatibility with our Golden Gate cloning system and cloned into the Level 0 CDS(ns) pICSL01005 acceptor. Once domesticated, the position of the Pikm HMA domain was substituted with pre-domesticated iGEM amilCP negative selection reporter cassettes internally flanked by outward pointing *Esp3I* sites to produce CAGA (5’) and GATG (3’) cloning overhangs. *Esp3I* was used to incorporate an iGEM RFP negative selection reporter cassette, internally flanked by outward pointing *BbsI* sites presenting CAGA (5’) and GATG (3’) cloning overhangs, allowing cloning of new domains via *BbsI* into the Pikm-1 DOM2 acceptor in the analogous position to where the HMA domain was located.

The RGA5 HMA and APB mutant were cloned into the Pikm-1 DOM2 acceptor via Golden Gate cloning with *BbsI* to assemble a full length Pikm-1 receptor chimera. Full length Pikm-1^RGA5^ and Pikm-1^APB^ were subsequently cloned into pICH47742 via *BsaI*, with a C-terminal 6xHIS/3xFLAG tag.

RGA5/RGA4 along with P19 were assembled into the binary agrobacterium expression vector pICSL4723 (Engler et al., 2014) via *BbsI*. RGA4 was tagged with a C-terminal 6xHA tag and RGA5 was left untagged to prevent effects on receptor function. Expression of RGA4 and RGA5 was driven by the *A. thaliana* actin and 2×35S promoters, respectively. For cell death assays, AVR-Pia was cloned untagged into pJK268c with P19, with expression driven by a 2×35S promoter. For co-IP assays, an N-terminally 4xMYC tagged AVR-Pia was cloned into pICH47752.

AVR-Pik effector variants used in this study were described previously (De la Concepcion et al., 2018). PWL2 was cloned into pICH47751 under a Ubi10 promoter and 35S terminator and C-terminal 4xMYC tag via Golden Gate cloning.

### Gene cloning – recombinant expression in *E. coli*

The RGA5, APB and Pikm-1 HMA domains as well as the AVR-Pik effector variants and AVR-Pia effector were cloned into pOPIN-GG vector pPGN-C (Bentham et al., 2021) with a cleavable N-terminal 6xHIS-GB1-3C tag via Golden Gate cloning with *BsaI*. AVR-Pii effector domain was cloned with a cleavable N-terminal MBP tag and an uncleavable C-terminal 6xHIS via In-fusion cloning into pOPINE (Berrow et al., 2007). For co-expression with the APB HMA for crystallography studies, AVR-PikF was cloned into pPGC-K (Bentham et al., 2021) without a tag via Golden Gate cloning with *BsaI*.

### In planta co-immunoprecipitation (co-IP)

Transient gene expression in planta was performed by infiltrating 4 week old *N. benthamiana* plants with *A. tumefaciens* strain GV3101 (C58 (rifR) Ti pMP90 (pTiC58DT-DNA) (gentR) Nopaline (pSoup-tetR)), grown at 22-25°C with high light intensity. *A. tumefaciens* carrying NLRs and effectors were infiltrated at OD_600_ 0.4 and 0.6, respectively, in agroinfiltration medium (10 mM MgCl_2_, 10 mM 2-(N-morpholine)-ethanesulfonic acid (MES), pH 5.6) with the addition of 150 µM acetosyringone.

Leaf tissue was collected 3 days post infiltrations (dpi) and frozen in liquid nitrogen before processing. Samples were ground to a fine powder in liquid nitrogen using a mortar and pestle before being mixed with two times weight/volume ice-cold Co-IP extraction buffer (25 mM Tris pH 7.5, 150 mM NaCl, 1 mM EDTA, 10 % glycerol, 2 % w/v PVPP, 10 mM DTT, 1 x cOmplete protease inhibitor tablet per 50 mL (Roche), 0.1 % Tween20). Samples were centrifuged at 4200 *g* at 4°C for 20 min, and supernatant was passed through a 0.45 µm Ministart syringe filter. SDS-PAGE/Western blot analysis was used to identify proteins in the sample with use of anti-FLAG M2 antibody (Sigma) and anti-MYC antibody (Santa Cruz Biotechnology) for NLRs and effectors, respectively.

For immunoprecipitation, 2 mL of filtered plant extract was incubated with 30 µL of M2 anti-FLAG magnetic beads (Sigma) in a rotary mixer for 3 hrs at 4°C. The FLAG beads were separated from the supernatant with use of a magnetic rack to allow for the removal of the supernatant. The beads were then washed with 1 mL of IP buffer (25 mM Tris pH 7.5, 150 mM NaCl, 1 mM EDTA, 10 % glycerol, 0.1 % Tween20). The FLAG beads were washed three times using this method. After washing, 30 µL of LDS Runblue sample buffer was added to the FLAG beads and incubated for 10 min at 70°C. The beads were then applied to a magnetic rack and the supernatant was loaded to SDS-PAGE gels and subsequently used for western blot analysis. PVDF membranes were probed with anti-FLAG M2 and anti-MYC antibodies to detect NLRs and effectors, respectively.

### *N. benthamiana* cell death assays and cell death scoring

Cell death assays and scoring were performed as described previously (De la Concepcion et al., 2021). In brief, *N. benthamiana* tissue was infiltrated with *A. tumefaciens* GV3101 (C58 [rifR] Ti pMP90 [pTiC58DT-DNA] [gentR] Nopaline [pSoup-tetR]) carrying NLRs and effectors at OD_600_ 0.4 and 0.6 respectively, and P19 at OD_600_ 0.1. Leaves were imaged 5 dpi from the abaxial side for UV fluorescence images. Images shown are representative of three independent experiments with internal technical repeats. The cell death scoring was performed using the cell death index previously presented in Maqbool et al., 2015. Dot plots were generated using R 4.0.5 (https://www.r-project.org) with the packages ggplot2 (Wickham, 2016). The size of the centre dot at each cell death value is directly proportional to the number of replicates in the sample with that score. All individual data points are represented as dots. Statistical analysis was performed using estimation graphics (Ho et al., 2019) with the besthr R package (De la Concepcion et al., 2021b; MacLean, 2019) and can be found in Appendix 1.

### Protein expression and purification from *E. coli*

Expression vectors containing the 6xHIS-GB1-tagged effectors and HMA domains were transformed into *E. coli* SHuffle cells. Using an overnight culture for inoculum, 8 L of SHuffle cells were grown in autoinduction media (AIM) at 30°C to an OD_600_ of 0.6 – 0.8 before the temperature was reduced to 18°C for overnight induction (Studier, 2005). Cells were pelleted by centrifugation at 5000 *g* for 10 mins and resuspended in lysis buffer (50 mM HEPES pH 8.0, 500 mM NaCl, 30 mM imidazole, 50 mM glycine and 5 % glycerol). Cell lysate was clarified by centrifugation at 45000 *g* for 20 mins following disruption of the resuspended pellet by sonication. Proteins were purified from clarified lysate via Ni^2+^ immobilised metal chromatography (IMAC) coupled with size-exclusion chromatography (SEC). The 6xHIS-GB1 tag was removed via overnight cleavage with 3C protease at 4°C before a final round of SEC using a buffer of 10 mM HEPES pH 8, 150 mM NaCl. Proteins were flash frozen in liquid nitrogen before storage at −80°C.

For co-expression of the APB/AVR-PikF complex, *E. coli* SHuffle cells were co-transformed with 6xHIS-GB1-tagged APB HMA and untagged AVR-PikF and plated on dual resistance carbenicillin and kanamycin selection. Expression and purification of the complex was then performed as described above, using dual selection for growth in large scale cultures.

### Crystallization, x-ray data collection, structure solution and refinement

The APB/AVR-PikF complex was concentrated to 10 mg/mL in SEC buffer (10 mM. HEPES pH 8.0, 150 mM NaCl) for crystallisation. Sitting drop, vapour diffusion crystallisation trials were set up in 96-well plants using an Oryx Nano robot (Douglas Instruments). Crystallisation plates were incubated at 20°C. APB/AVR-PikF crystals appeared in the SG1 ™ Screen (Molecular Dimensions) after 10 days in a 0.1 M BIS-TRIS pH 5.5, 25 % PEG 3350 condition. Crystals were harvested and snap frozen in liquid nitrogen prior to shipping.

Crystals of the APB/AVR-PikF complex diffracted to 1.3 Å and x-ray datasets were collected at the Diamond Light Source on the i04 beamline under proposal mx25108. The data were processed using the xia2 pipeline and AIMLESS as implemented in CCP4i2 (Winn et al., 2011). Using the structure of the OsHIPP19/AVR-PikF complex (PDB ID: 7B1I) as a template, the structure of the APB/AVR-PikF complex was solved using molecular replacement with PHASER (McCoy et al., 2007). The final structure was obtained after iterative cycles of refinement using COOT and REFMAC (Emsley and Cowtan, 2004; Murshudov et al., 1997). Structure geometry was validated using the tools in COOT and MOLPROBITY (Chen et al., 2010; Emsley and Cowtan, 2004). Protein interface analyses were performed using QtPISA and ChimeraX (Krissinel and Henrick, 2007; Pettersen et al., 2021). Models are visualised using ChimeraX (Pettersen et al., 2021). X-ray diffraction data can be found in the Protein Data Bank (https://www.ebi.ac.uk/pdbe/) under the accession number 8B2R.

### Analytical size-exclusion chromatography

150 µg of purified AVR-PikF was mixed with 150 µg of the RGA5 and APB HMA domains and incubated on ice for 30 mins before separation via SEC using a Superdex S75 10/300 GL size-exclusion column (Cytiva). As a negative control, 150 µg of AVR-PikF was run alone. HMA domains were not run separate from AVR-PikF due to low or no absorbance at A_280_ resulting in no observable peak in the chromatogram. Chromatograms were visualised using the ggplot2 R library in R 4.0.5 (Wickham, 2016).

### Biophysical analysis with surface plasmon resonance

Surface plasmon resonance was performed using a Biacore 8K (Cytiva). Purified HMA domains were immobilised on a Series S Sensor CM5 Chip (Cytiva) via amine-coupling using 0.4 M 1-ethyl-3-(3-dimethylaminopropyl)-carbodiimide (EDC) and 0.1 M N-hydroxysuccinimide (NHS) to activate the chip surface prior to binding of HMAs at two concentrations on different channels, a high concentration (30 nM, ∼2000 response units (RU)) and a low concentration (0.3 nM, ∼200 RU) to allow for accurate measurement of affinity and kinetics of strong and weak interactors. 1 M ethanolamine-HCl pH 8.5 was then used to block the CM5 chip after coupling was completed.

Samples were run in HBS-EP+ running buffer (0.1 M HEPES, 1.5 M NaCl, 0.03 M EDTA and 0.5% v/v Tween20) and the chip was regenerated after each cycle with an ionic regeneration buffer (0.46 M KSCN, 1.83 M MgCl_2_, 0.92 M urea, 1.83 M guanidine-HCl). Effectors were run over the chip at a flow rate of 100 µL/min; contact and dissociation time varied depending on the experiment (see below).

Where possible, we performed multicycle kinetics to assess the affinity and binding kinetics of the effectors for the HMA. For strong interactions (Pikm-1 HMA with AVR-PikD) we used serial dilutions of effectors from 50 nM – 0 nM, and for weak interactions we used serial dilutions of 50 µM – 0 µM (AVR-Pia with RGA5 HMA, APB HMA and Pikm-1 HMA; AVR-PikC with RGA5 HMA and Pikm-1 HMA; AVR-PikF with RGA5 HMA and Pikm-1 HMA), with each concentration being performed in triplicate. Contact time and dissociation times for the experiment were set at 120 s.

For strong interactions we performed single-cycle kinetics due to the extremely slow dissociation rates of the effectors from the HMA domains, which interfered with accurate calculations of kinetic parameters and binding affinity. For single cycle kinetics, increasing concentrations of effector (0 nM – 50 nM) were sequentially flowed over the HMA-bound sensor chip each with a contact time of 120 s before a single dissociation phase of 600 s. Each cycle was performed in triplicate.

SPR sensograms were analysed with the Biacore Insight Evaluation Software (Cytiva) and equilibrium dissociation constants (*K*_D_) values were calculated using a 1:1 binding model from a kinetic fit model. Residual graphs are generated from the subtraction of the experimental data from the fit model (ΔFit – Exp). Sensograms and residual graphs were generated in R 4.0.5 using the ggplot2 R package (Wickham, 2016).

### Appendix 1. Statistical analysis of cell death scoring with besthr.

Cell death scoring in this study was performed through qualitative measurement of cell death as determined by approximation of autofluorescence under UV light 5 dpi, as previously performed in De la Concepcion et al., 2019. The autofluorescence was compared to a previously established cell death scale (Maqbool et al., 2015). To analyse our cell death scoring, we used estimation methods (Ho et al., 2019) and visualised these with use of the besthr R package (MacLean, 2019) to generate estimation graphics.

Besthr compares the cell death scores of all samples and ranks them irrespective of sample then generates mean ranks for the control and test samples. A bootstrap process is then performed on the ranked test data in which samples of equal size to the experiment were replaced and a mean rank is calculated.

Rank means were calculated after 1000 bootstrap samples and a distribution of the mean ranks were plotted, with the 2.5 and 97.5 quantiles calculated and highlighted on the plotted distribution. If the mean of the control data is outside of the 2.5 or 97.5 quantile boundaries, the control and test means are considered to be different.

---

**Appendix 1 A.**
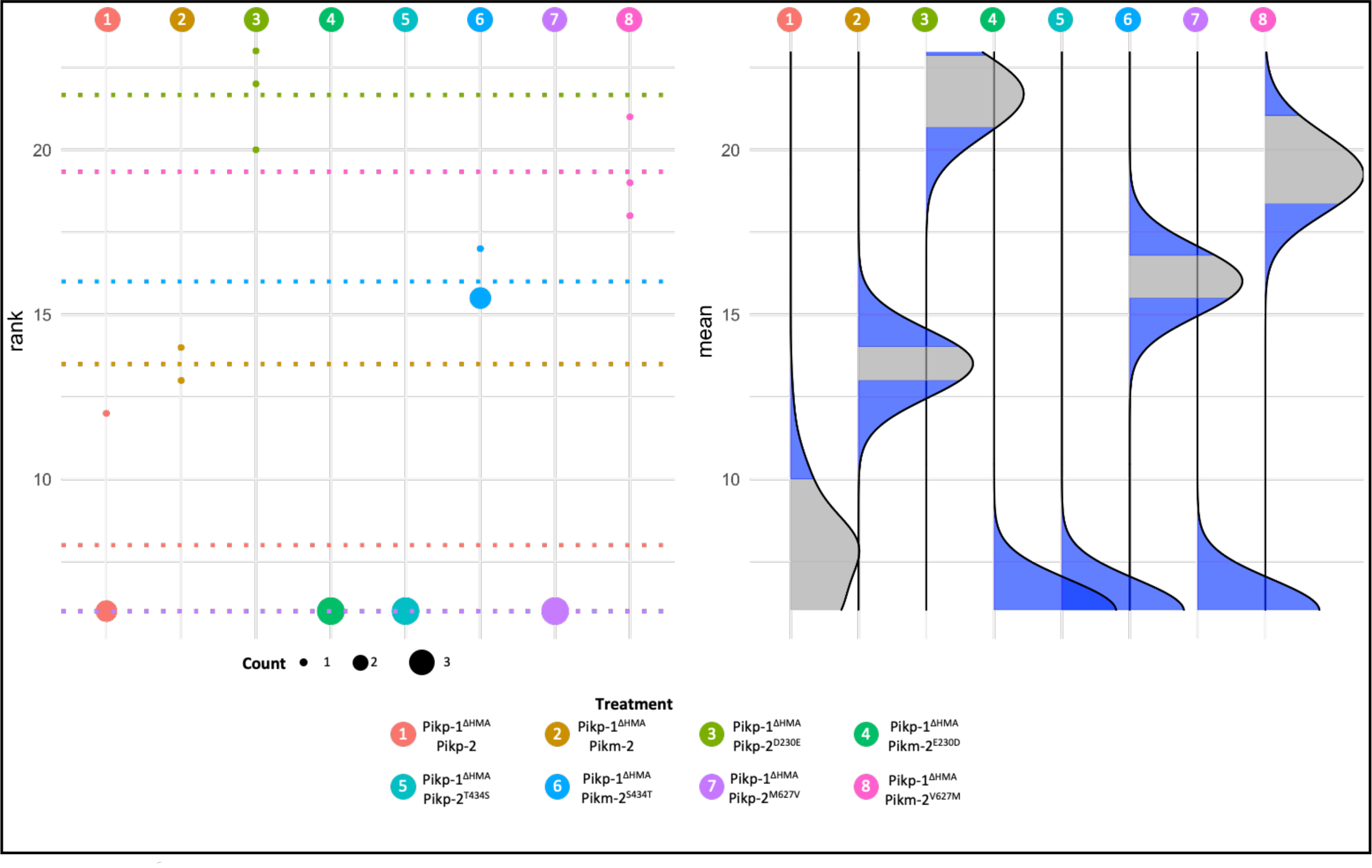
Statistical analysis of cell death scoring from Fig 1.

**Appendix 1 B.**
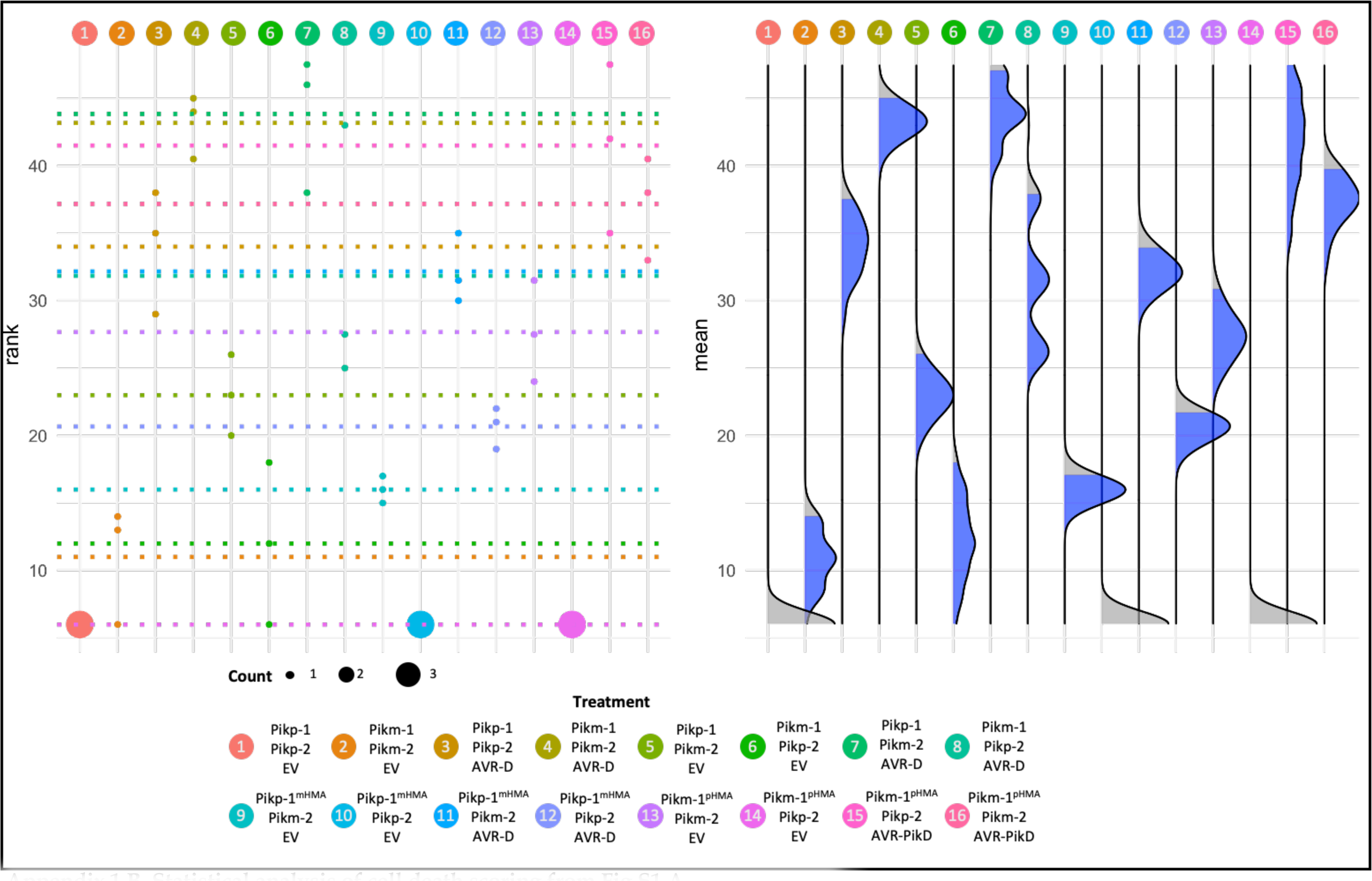
Statistical analysis of cell death scoring from Fig S1 A.

**Appendix 1 C.**
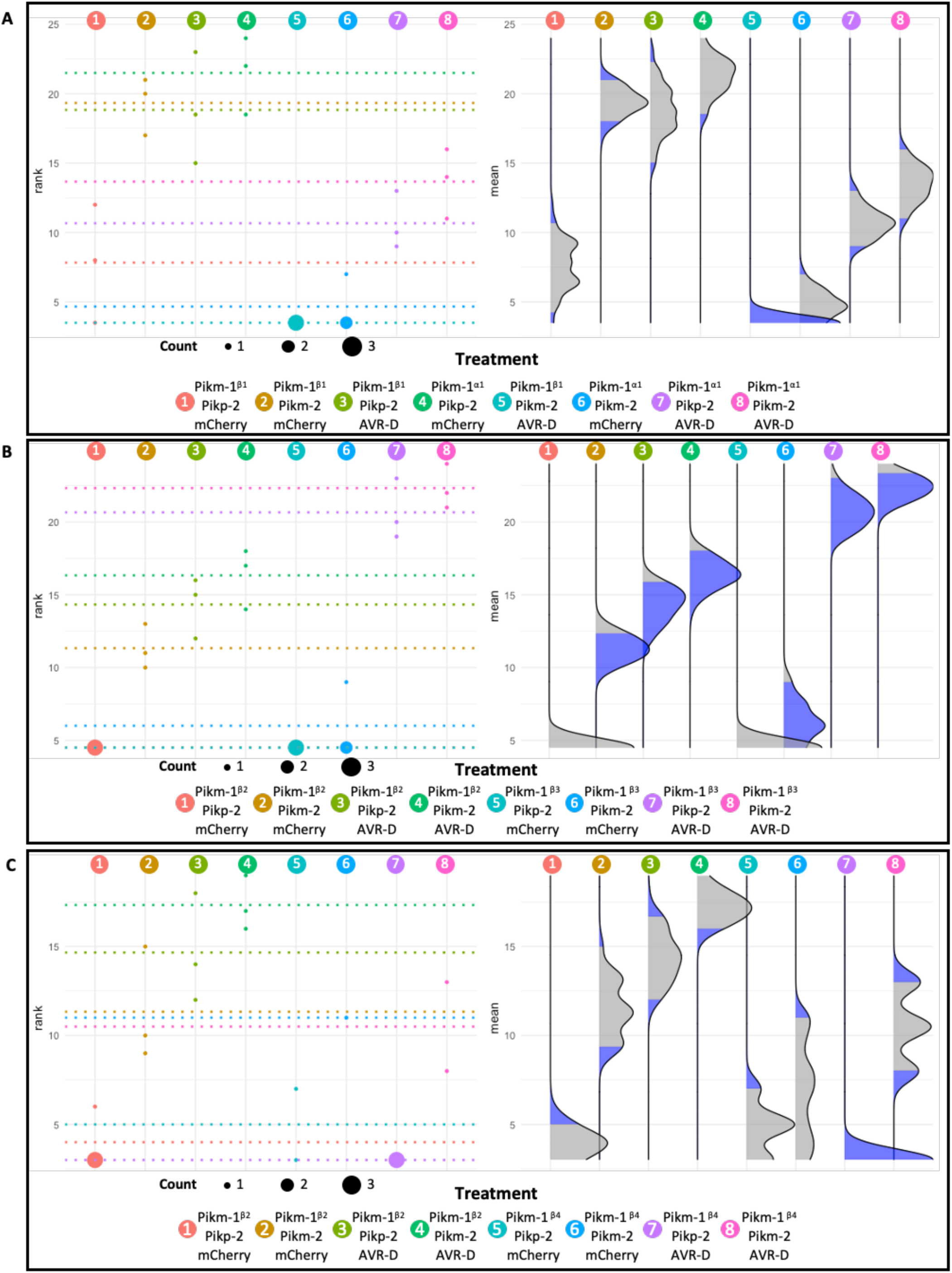
Statistical analysis of cell death scoring from Fig S1 B. **A)** Chimeras of Pikm-1 with the β1 and α1 secondary structures of Pikp-1 HMA **B)** Chimeras of Pikm-1 with the β2 and β3 structures of Pikp-1 **C)** Chimeras of Pikm-1 with the α2 and β4 secondary structures of Pikp-1 HMA

**Appendix 1 D.**
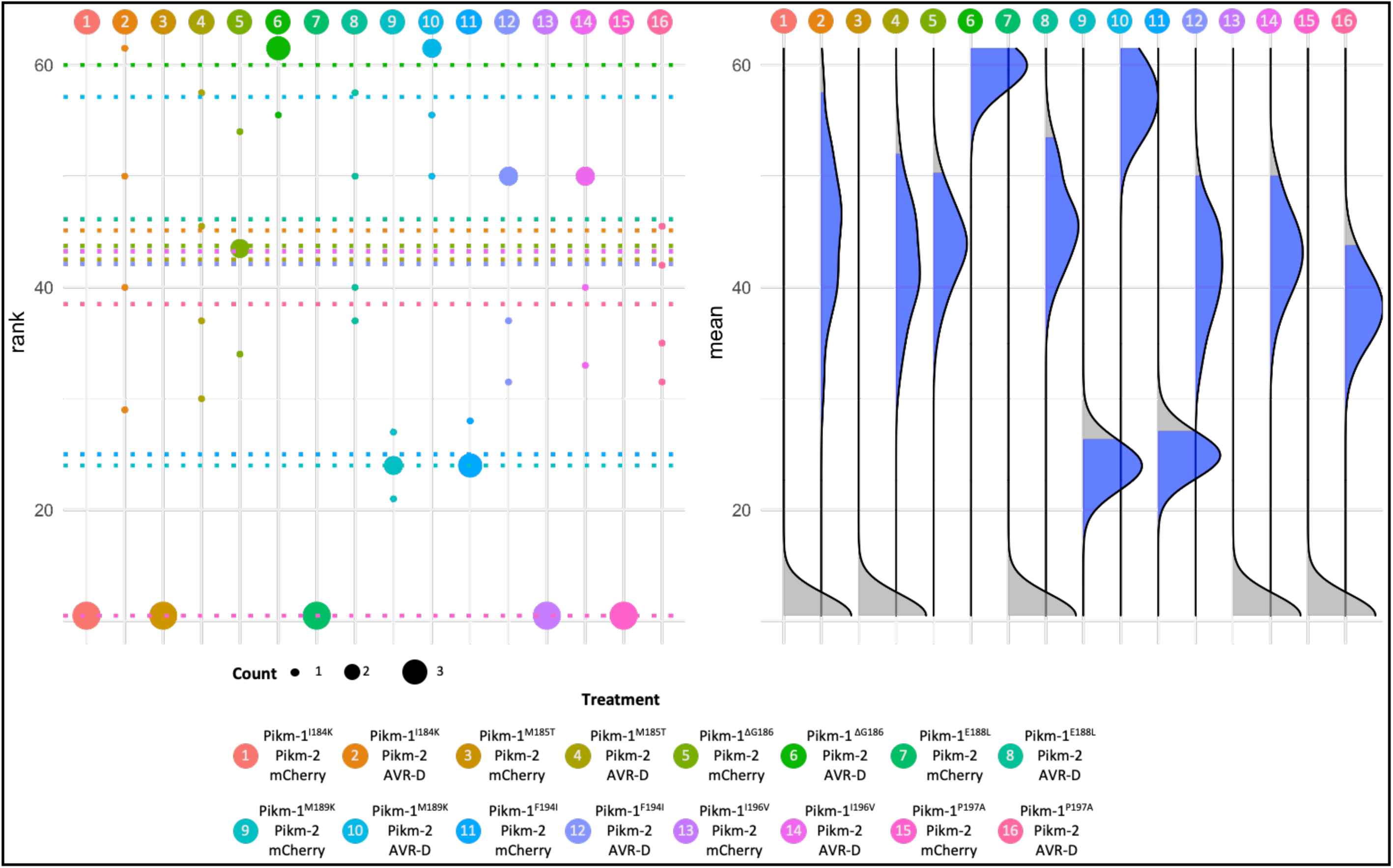
Statistical analysis of cell death scoring from Fig S3 A.

**Appendix 1 E.**
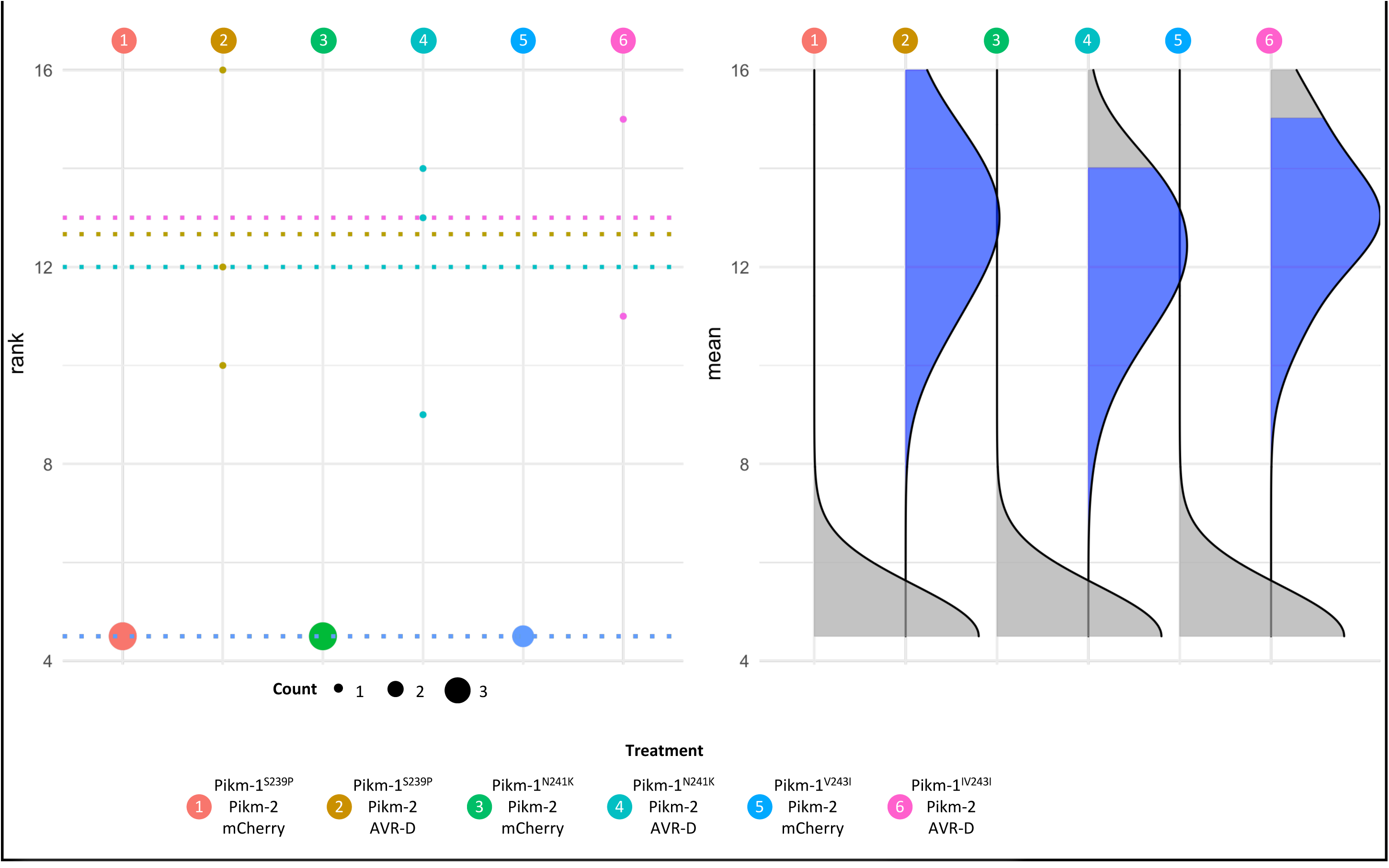
Statistical analysis of cell death scoring from Fig S3 B.

**Appendix 1 F.**
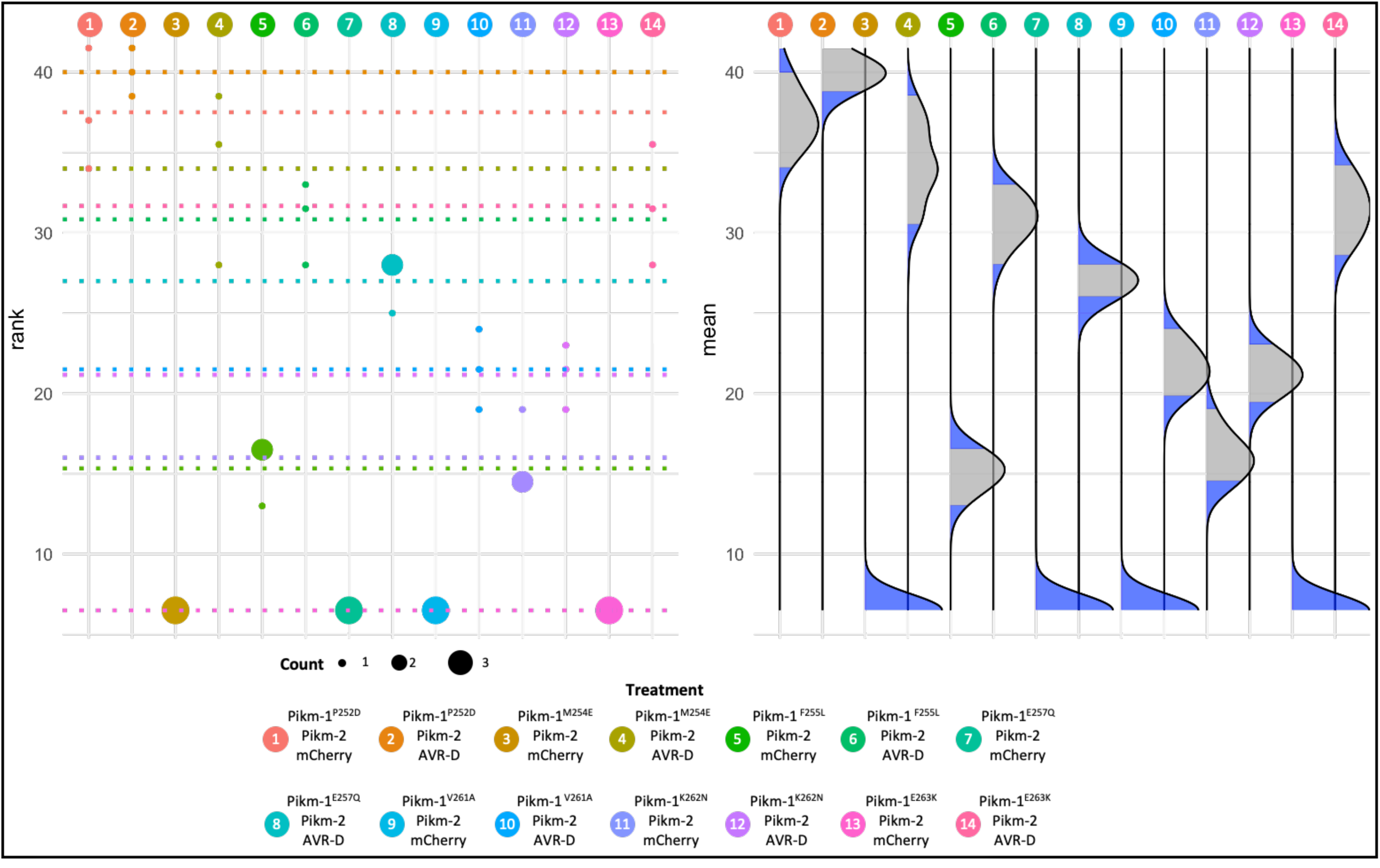
Statistical analysis of cell death scoring from Fig S3 C.

**Appendix 1 G.**
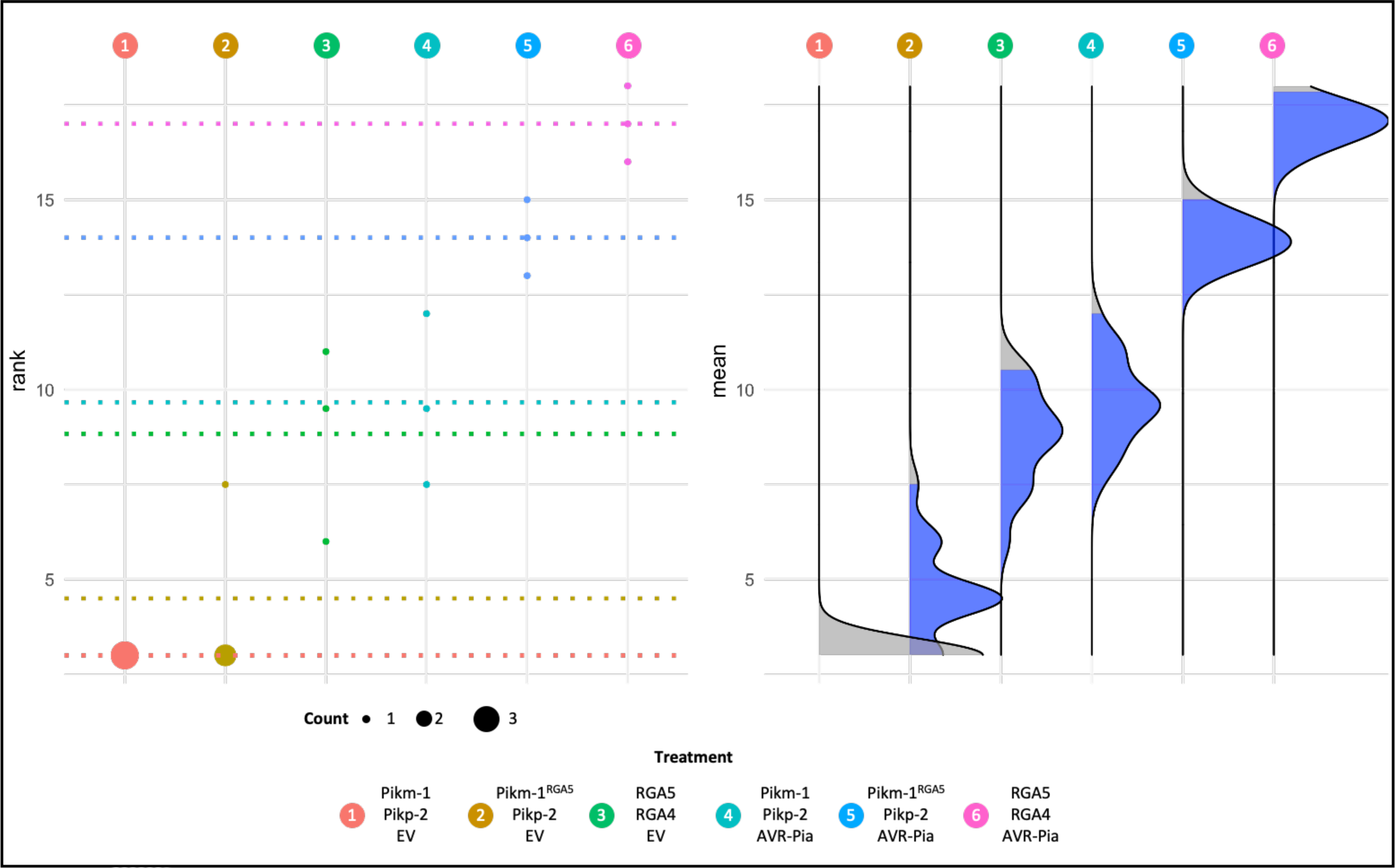
Statistical analysis of cell death scoring from Fig 3 B.

**Appendix 1 H.**
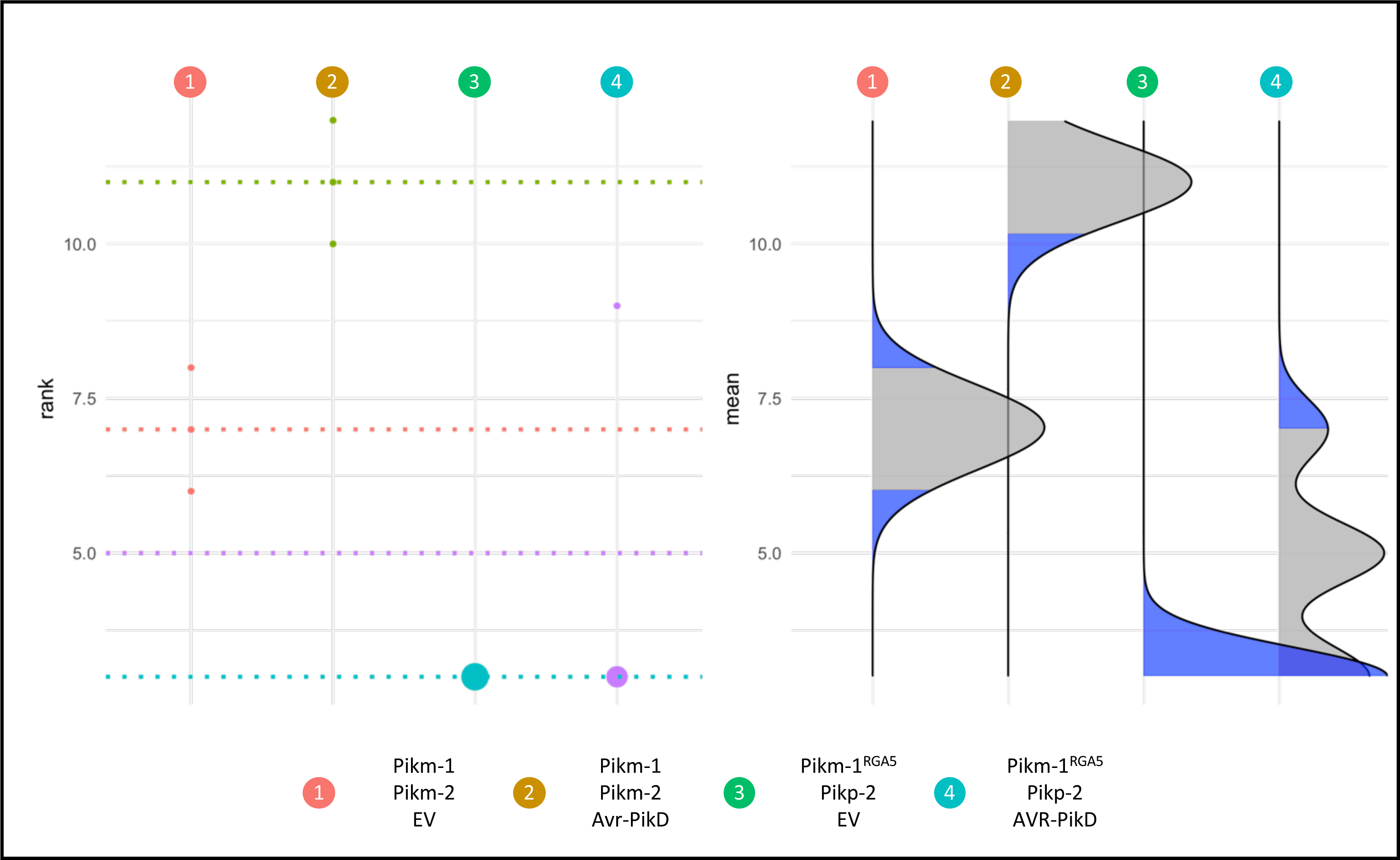
Statistical analysis of cell death scoring from Fig S4.

**Appendix 1 I.**
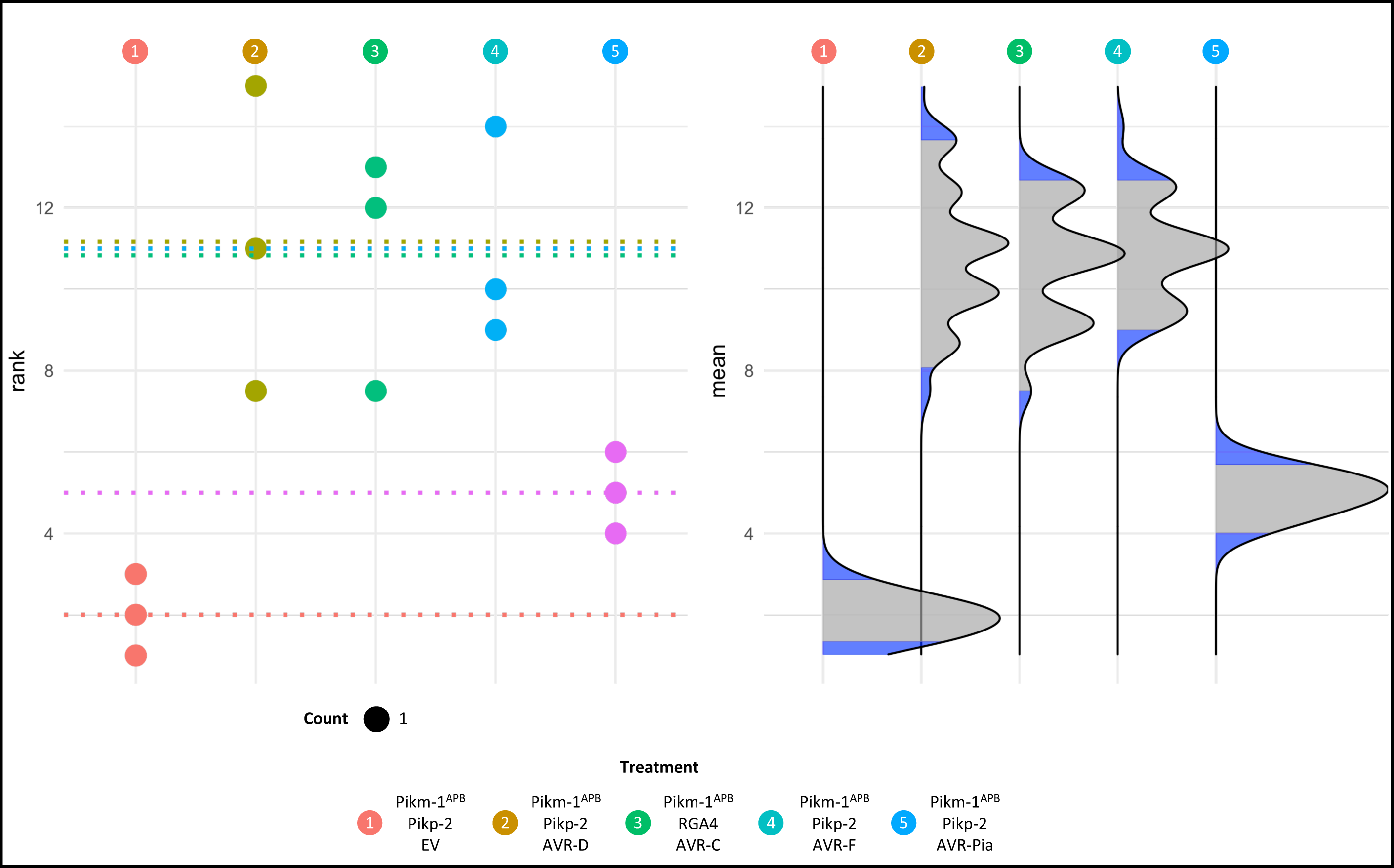
Statistical analysis of cell death scoring from Fig 4 B.

## References

1. Adachi, H., Derevnina, L., Kamoun, S., 2019. NLR singletons, pairs, and networks: evolution, assembly, and regulation of the intracellular immunoreceptor circuitry of plants. Curr Opin Plant Biol 50, 121–131. https://doi.org/10.1016/j.pbi.2019.04.007

2. Baggs, E., Dagdas, G., Krasileva, K., 2017. NLR diversity, helpers and integrated domains: making sense of the NLR IDentity. Current Opinion in Plant Biology, 38 Biotic interactions 2017 38, 59–67. https://doi.org/10.1016/j.pbi.2017.04.012

3. Bentham, A.R., De la Concepcion, J.C., Mukhi, N., Zdrzałek, R., Draeger, M., Gorenkin, D., Hughes, R.K., Banfield, M.J., 2020. A molecular roadmap to the plant immune system. JBC 295, 14916– 14935. https://doi.org/10.1074/jbc.REV120.010852

4. Bentham, A.R., Youles, M., Mendel, M.N., Varden, F.A., Concepcion, J.C.D. la, Banfield, M.J., 2021. pOPIN-GG: A resource for modular assembly in protein expression vectors. https://doi.org/10.1101/2021.08.10.455798

5. Berrow, N.S., Alderton, D., Sainsbury, S., Nettleship, J., Assenberg, R., Rahman, N., Stuart, D.I., Owens, R.J., 2007. A versatile ligation-independent cloning method suitable for high-throughput expression screening applications. Nucleic Acids Res 35, e45. https://doi.org/10.1093/nar/gkm047

6. Białas, A., Langner, T., Harant, A., Contreras, M.P., Stevenson, C.E., Lawson, D.M., Sklenar, J., Kellner, R., Moscou, M.J., Terauchi, R., Banfield, M.J., Kamoun, S., 2021. Two NLR immune receptors acquired high-affinity binding to a fungal effector through convergent evolution of their integrated domain. eLife 10, e66961. https://doi.org/10.7554/eLife.66961

7. Białas, A., Zess, E.K., De la Concepcion, J.C., Franceschetti, M., Pennington, H.G., Yoshida, K., Upson, J.L., Chanclud, E., Wu, C.-H., Langner, T., Maqbool, A., Varden, F.A., Derevnina, L., Belhaj, K., Fujisaki, K., Saitoh, H., Terauchi, R., Banfield, M.J., Kamoun, S., 2018. Lessons in Effector and NLR Biology of Plant-Microbe Systems. MPMI 31, 34–45. https://doi.org/10.1094/MPMI-08-17-0196-FI

8. Bomblies, K., Lempe, J., Epple, P., Warthmann, N., Lanz, C., Dangl, J.L., Weigel, D., 2007. Autoimmune response as a mechanism for a Dobzhansky-Muller-type incompatibility syndrome in plants. PLoS Biol 5, e236. https://doi.org/10.1371/journal.pbio.0050236

9. Burdett, H., Kobe, B., Anderson, P.A., 2019. Animal NLRs continue to inform plant NLR structure and function. Archives of Biochemistry and Biophysics, Inflammasomes: Intracellular mediators of immune defence 670, 58–68. https://doi.org/10.1016/j.abb.2019.05.001

10. Calvo-Baltanás, V., Wang, J., Chae, E., 2021. Hybrid Incompatibility of the Plant Immune System: An Opposite Force to Heterosis Equilibrating Hybrid Performances. Frontiers in Plant Science 11.

11. Cesari, S., 2018. Multiple strategies for pathogen perception by plant immune receptors. New Phytologist 219, 17–24. https://doi.org/10.1111/nph.14877

12. Cesari, S., Bernoux, M., Moncuquet, P., Kroj, T., Dodds, P.N., 2014. A novel conserved mechanism for plant NLR protein pairs: the “integrated decoy” hypothesis. Front Plant Sci 5, 606. https://doi.org/10.3389/fpls.2014.00606

13. Cesari, S., Thilliez, G., Ribot, C., Chalvon, V., Michel, C., Jauneau, A., Rivas, S., Alaux, L., Kanzaki, H., Okuyama, Y., Morel, J.-B., Fournier, E., Tharreau, D., Terauchi, R., Kroj, T., 2013. The Rice Resistance Protein Pair RGA4/RGA5 Recognizes the Magnaporthe oryzae Effectors AVR-Pia and AVR1-CO39 by Direct Binding. The Plant Cell 25, 1463–1481. https://doi.org/10.1105/tpc.112.107201

14. Cesari, S., Xi, Y., Declerck, N., Chalvon, V., Mammri, L., Pugnière, M., Henriquet, C., de Guillen, K., Chochois, V., Padilla, A., Kroj, T., 2022. New recognition specificity in a plant immune receptor by molecular engineering of its integrated domain. Nat Commun 13, 1524. https://doi.org/10.1038/s41467-022-29196-6

15. Chae, E., Bomblies, K., Kim, S.-T., Karelina, D., Zaidem, M., Ossowski, S., Martín-Pizarro, C., Laitinen, R.A.E., Rowan, B.A., Tenenboim, H., Lechner, S., Demar, M., Habring-Müller, A., Lanz, C., Rätsch, G., Weigel, D., 2014. Species-wide genetic incompatibility analysis identifies immune genes as hot spots of deleterious epistasis. Cell 159, 1341–1351. https://doi.org/10.1016/j.cell.2014.10.049

16. Chen, V.B., Arendall, W.B., Headd, J.J., Keedy, D.A., Immormino, R.M., Kapral, G.J., Murray, L.W., Richardson, J.S., Richardson, D.C., 2010. MolProbity: all-atom structure validation for macromolecular crystallography. Acta Crystallogr D Biol Crystallogr 66, 12–21. https://doi.org/10.1107/S0907444909042073

17. Costanzo, S., Jia, Y., 2010. Sequence variation at the rice blast resistance gene Pi-km locus: Implications for the development of allele specific markers. Plant Science 178, 523–530. https://doi.org/10.1016/j.plantsci.2010.02.014

18. De la Concepcion, J.C., Franceschetti, M., MacLean, D., Terauchi, R., Kamoun, S., Banfield, M.J., 2019. Protein engineering expands the effector recognition profile of a rice NLR immune receptor. eLife 8, e47713. https://doi.org/10.7554/eLife.47713

19. De la Concepcion, J.C., Franceschetti, M., Maqbool, A., Saitoh, H., Terauchi, R., Kamoun, S., Banfield, M.J., 2018. Polymorphic residues in rice NLRs expand binding and response to effectors of the blast pathogen. Nature Plants 4, 576–585. https://doi.org/10.1038/s41477-018-0194-x

20. De la Concepcion, J.C., Fujisaki, K., Bentham, A.R., Mireles, N.C., Hernandez, V.S. de M., Shimizu, M., Lawson, D.M., Kamoun, S., Terauchi, R., Banfield, M.J., 2022. Binding of a blast fungus Zinc-finger fold effector to a hydrophobic pocket in the host exocyst subunit Exo70 modulates immune recognition in rice. https://doi.org/10.1101/2022.06.18.496527

21. De la Concepcion, J.C., Maidment, J.H.R., Longya, A., Xiao, G., Franceschetti, M., Banfield, M.J., 2021a. The allelic rice immune receptor Pikh confers extended resistance to strains of the blast fungus through a single polymorphism in the effector binding interface. PLOS Pathogens 17, e1009368. https://doi.org/10.1371/journal.ppat.1009368

22. De la Concepcion, J.C., Vega Benjumea, J., Bialas, A., Terauchi, R., Kamoun, S., Banfield, M.J., 2021b. Functional diversification gave rise to allelic specialization in a rice NLR immune receptor pair. eLife 10, e71662. https://doi.org/10.7554/eLife.71662

23. Emsley, P., Cowtan, K., 2004. Coot: model-building tools for molecular graphics. Acta Crystallogr D Biol Crystallogr 60, 2126–2132. https://doi.org/10.1107/S0907444904019158

24. Engler, C., Youles, M., Gruetzner, R., Ehnert, T.-M., Werner, S., Jones, J.D.G., Patron, N.J., Marillonnet, S., 2014. A Golden Gate Modular Cloning Toolbox for Plants. ACS Synth. Biol. 3, 839–843. https://doi.org/10.1021/sb4001504

25. Feehan, J.M., Castel, B., Bentham, A.R., Jones, J.D., 2020. Plant NLRs get by with a little help from their friends. Curr Opin Plant Biol 56, 99–108. https://doi.org/10.1016/j.pbi.2020.04.006

26. Fujisaki, K., Abe, Y., Kanzaki, E., Ito, K., Utsushi, H., Saitoh, H., Białas, A., Banfield, M.J., Kamoun, S., Terauchi, R., 2017. An unconventional NOI/RIN4 domain of a rice NLR protein binds host EXO70 protein to confer fungal immunity. https://doi.org/10.1101/239400

27. Guo, L., Cesari, S., de Guillen, K., Chalvon, V., Mammri, L., Ma, M., Meusnier, I., Bonnot, F., Padilla, A., Peng, Y.-L., Liu, J., Kroj, T., 2018. Specific recognition of two MAX effectors by integrated HMA domains in plant immune receptors involves distinct binding surfaces. Proc. Natl. Acad. Sci. U.S.A. 115, 11637–11642. https://doi.org/10.1073/pnas.1810705115

28. Ho, J., Tumkaya, T., Aryal, S., Choi, H., Claridge-Chang, A., 2019. Moving beyond P values: data analysis with estimation graphics. Nat Methods 16, 565–566. https://doi.org/10.1038/s41592-019-0470-3

29. Jones, J.D.G., Vance, R.E., Dangl, J.L., 2016. Intracellular innate immune surveillance devices in plants and animals. Science 354, aaf6395. https://doi.org/10.1126/science.aaf6395

30. Jumper, J., Evans, R., Pritzel, A., Green, T., Figurnov, M., Ronneberger, O., Tunyasuvunakool, K., Bates, R., Žídek, A., Potapenko, A., Bridgland, A., Meyer, C., Kohl, S.A.A., Ballard, A.J., Cowie, A., Romera-Paredes, B., Nikolov, S., Jain, R., Adler, J., Back, T., Petersen, S., Reiman, D., Clancy, E., Zielinski, M., Steinegger, M., Pacholska, M., Berghammer, T., Bodenstein, S., Silver, D., Vinyals, O., Senior, A.W., Kavukcuoglu, K., Kohli, P., Hassabis, D., 2021. Highly accurate protein structure prediction with AlphaFold. Nature 596, 583–589. https://doi.org/10.1038/s41586-021-03819-2

31. Kanzaki, H., Yoshida, K., Saitoh, H., Fujisaki, K., Hirabuchi, A., Alaux, L., Fournier, E., Tharreau, D., Terauchi, R., 2012. Arms race co-evolution of Magnaporthe oryzae AVR-Pik and rice Pik genes driven by their physical interactions. Plant J 72, 894–907. https://doi.org/10.1111/j.1365-313X.2012.05110.x

32. Kourelis, J., Adachi, H., 2022. Activation and Regulation of NLR Immune Receptor Networks. Plant and Cell Physiology pcac116. https://doi.org/10.1093/pcp/pcac116

33. Kourelis, J., Marchal, C., Kamoun, S., 2021. NLR immune receptor-nanobody fusions confer plant disease resistance. https://doi.org/10.1101/2021.10.24.465418

34. Krissinel, E., Henrick, K., 2007. Inference of macromolecular assemblies from crystalline state. J Mol Biol 372, 774–797. https://doi.org/10.1016/j.jmb.2007.05.022

35. Kroj, T., Chanclud, E., Michel-Romiti, C., Grand, X., Morel, J.-B., 2016. Integration of decoy domains derived from protein targets of pathogen effectors into plant immune receptors is widespread. New Phytologist 210, 618–626. https://doi.org/10.1111/nph.13869

36. Le Roux, C., Huet, G., Jauneau, A., Camborde, L., Trémousaygue, D., Kraut, A., Zhou, B., Levaillant, M., Adachi, H., Yoshioka, H., Raffaele, S., Berthomé, R., Couté, Y., Parker, J.E., Deslandes, L., 2015. A Receptor Pair with an Integrated Decoy Converts Pathogen Disabling of Transcription Factors to Immunity. Cell 161, 1074–1088. https://doi.org/10.1016/j.cell.2015.04.025

37. Liu, Y., Zhang, X., Yuan, G., Wang, D., Zheng, Y., Ma, M., Guo, L., Bhadauria, V., Peng, Y.-L., Liu, J., 2021. A designer rice NLR immune receptor confers resistance to the rice blast fungus carrying noncorresponding avirulence effectors. Proceedings of the National Academy of Sciences 118, e2110751118. https://doi.org/10.1073/pnas.2110751118

38. Lüdke, D., Yan, Q., Rohmann, P.F.W., Wiermer, M., 2022. NLR we there yet? Nucleocytoplasmic coordination of NLR-mediated immunity. New Phytologist n/a. https://doi.org/10.1111/nph.18359

39. MacLean, D., 2019. TeamMacLean/besthr: Initial Release. https://doi.org/10.5281/zenodo.3374507

40. Maidment, J.H.R., Franceschetti, M., Maqbool, A., Saitoh, H., Jantasuriyarat, C., Kamoun, S., Terauchi, R., Banfield, M.J., 2021. Multiple variants of the fungal effector AVR-Pik bind the HMA domain of the rice protein OsHIPP19, providing a foundation to engineer plant defense. Journal of Biological Chemistry 296. https://doi.org/10.1016/j.jbc.2021.100371

41. Maidment, J.H.R., Shimizu, M., Vera, S., Franceschetti, M., Longya, A., Stevenson, C.E.M., Concepcion, J.D. la, Białas, A., Kamoun, S., Terauchi, R., Banfield, M.J., 2022. Effector target-guided engineering of an integrated domain expands the disease resistance profile of a rice NLR immune receptor. https://doi.org/10.1101/2022.06.14.496076

42. Maqbool, A., Saitoh, H., Franceschetti, M., Stevenson, C., Uemura, A., Kanzaki, H., Kamoun, S., Terauchi, R., Banfield, M., 2015. Structural basis of pathogen recognition by an integrated HMA domain in a plant NLR immune receptor. eLife 4, e08709. https://doi.org/10.7554/eLife.08709

43. Marchal, C., Michalopoulou, V.A., Zou, Z., Cevik, V., Sarris, P.F., 2022. Show me your ID: NLR immune receptors with integrated domains in plants. Essays in Biochemistry EBC20210084. https://doi.org/10.1042/EBC20210084

44. Maruta, N., Burdett, H., Lim, B.Y.J., Hu, X., Desa, S., Manik, M.K., Kobe, B., 2022. Structural basis of NLR activation and innate immune signalling in plants. Immunogenetics 74, 5–26. https://doi.org/10.1007/s00251-021-01242-5

45. McCoy, A.J., Grosse-Kunstleve, R.W., Adams, P.D., Winn, M.D., Storoni, L.C., Read, R.J., 2007. Phaser crystallographic software. J Appl Crystallogr 40, 658–674. https://doi.org/10.1107/S0021889807021206

46. Mirdita, M., Schütze, K., Moriwaki, Y., Heo, L., Ovchinnikov, S., Steinegger, M., 2022. ColabFold: making protein folding accessible to all. Nat Methods 19, 679–682. https://doi.org/10.1038/s41592-022-01488-1

47. Monteiro, F., Nishimura, M.T., 2018. Structural, Functional, and Genomic Diversity of Plant NLR Proteins: An Evolved Resource for Rational Engineering of Plant Immunity. Annu. Rev. Phytopathol. 56, 243–267. https://doi.org/10.1146/annurev-phyto-080417-045817

48. Mukhi, N., Brown, H., Gorenkin, D., Ding, P., Bentham, A.R., Jones, J.D.G., Banfield, M.J., 2021. Perception of structurally distinct effectors by the integrated WRKY domain of a plant immune receptor. https://doi.org/10.1101/2021.07.28.454147

49. Murshudov, G.N., Vagin, A.A., Dodson, E.J., 1997. Refinement of macromolecular structures by the maximum-likelihood method. Acta Crystallogr D Biol Crystallogr 53, 240–255. https://doi.org/10.1107/S0907444996012255

50. Ordon, J., Martin, P., Erickson, J.L., Ferik, F., Balcke, G., Bonas, U., Stuttmann, J., 2021. Disentangling cause and consequence: genetic dissection of the DANGEROUS MIX2 risk locus, and activation of the DM2h NLR in autoimmunity. The Plant Journal 106, 1008–1023. https://doi.org/10.1111/tpj.15215

51. Ortiz, D., de Guillen, K., Cesari, S., Chalvon, V., Gracy, J., Padilla, A., Kroj, T., 2017. Recognition of the Magnaporthe oryzae Effector AVR-Pia by the Decoy Domain of the Rice NLR Immune Receptor RGA5[OPEN]. Plant Cell 29, 156–168. https://doi.org/10.1105/tpc.16.00435

52. Outram, M.A., Figueroa, M., Sperschneider, J., Williams, S.J., Dodds, P.N., 2022. Seeing is believing: Exploiting advances in structural biology to understand and engineer plant immunity. Current Opinion in Plant Biology 67, 102210. https://doi.org/10.1016/j.pbi.2022.102210

53. Pettersen, E.F., Goddard, T.D., Huang, C.C., Meng, E.C., Couch, G.S., Croll, T.I., Morris, J.H., Ferrin, T.E., 2021. UCSF ChimeraX: Structure visualization for researchers, educators, and developers. Protein Sci 30, 70–82. https://doi.org/10.1002/pro.3943

54. Sarris, P.F., Cevik, V., Dagdas, G., Jones, J.D.G., Krasileva, K.V., 2016. Comparative analysis of plant immune receptor architectures uncovers host proteins likely targeted by pathogens. BMC Biology 14, 8. https://doi.org/10.1186/s12915-016-0228-7

55. Studier, F.W., 2005. Protein production by auto-induction in high-density shaking cultures. Protein Expression and Purification 41, 207–234. https://doi.org/10.1016/j.pep.2005.01.016

56. Sugihara, Y., Abe, Y., Takagi, H., Abe, A., Shimizu, M., Ito, K., Kanzaki, E., Oikawa, K., Kourelis, J., Langner, T., Win, J., Białas, A., Lüdke, D., Chuma, I., Saitoh, H., Kobayashi, M., Zheng, S., Tosa, Y., Banfield, M.J., Kamoun, S., Terauchi, R., Fujisaki, K., 2022. Tangled gene-for-gene interactions mediate co-evolution of the rice NLR immune receptor Pik and blast fungus effector proteins. https://doi.org/10.1101/2022.07.19.500555

57. Takken, F.L.W., Goverse, A., 2012. How to build a pathogen detector: structural basis of NB-LRR function. Curr Opin Plant Biol 15, 375–384. https://doi.org/10.1016/j.pbi.2012.05.001

58. Tamborski, J., Seong, K., Liu, F., Staskawicz, B., Krasileva, K.V., 2022. Engineering of Sr33 and Sr50 plant immune receptors to alter recognition specificity and autoactivity. https://doi.org/10.1101/2022.03.05.483131

59. Tran, D.T.N., Chung, E.-H., Habring-Müller, A., Demar, M., Schwab, R., Dangl, J.L., Weigel, D., Chae, E., 2017. Activation of a Plant NLR Complex through Heteromeric Association with an Autoimmune Risk Variant of Another NLR. Current Biology 27, 1148–1160. https://doi.org/10.1016/j.cub.2017.03.018

60. Varden, F.A., Saitoh, H., Yoshino, K., Franceschetti, M., Kamoun, S., Terauchi, R., Banfield, M.J., 2019. Cross-reactivity of a rice NLR immune receptor to distinct effectors from the rice blast pathogen Magnaporthe oryzae provides partial disease resistance. J Biol Chem 294, 13006– 13016. https://doi.org/10.1074/jbc.RA119.007730

61. Wickham, H., 2016. ggplot2: Elegant Graphics for Data Analysis. Springer-Verlag New York.

62. Winn, M.D., Ballard, C.C., Cowtan, K.D., Dodson, E.J., Emsley, P., Evans, P.R., Keegan, R.M., Krissinel, E.B., Leslie, A.G.W., McCoy, A., McNicholas, S.J., Murshudov, G.N., Pannu, N.S., Potterton, E.A., Powell, H.R., Read, R.J., Vagin, A., Wilson, K.S., 2011. Overview of the CCP4 suite and current developments. Acta Crystallogr D Biol Crystallogr 67, 235–242. https://doi.org/10.1107/S0907444910045749

63. Wu, C.-H., Derevnina, L., Kamoun, S., 2018. Receptor networks underpin plant immunity. Science 360, 1300–1301. https://doi.org/10.1126/science.aat2623

64. Zdrzałek, R., Kamoun, S., Terauchi, R., Saitoh, H., Banfield, M.J., 2020. The rice NLR pair Pikp-1/Pikp-2 initiates cell death through receptor cooperation rather than negative regulation. PLoS One 15, e0238616. https://doi.org/10.1371/journal.pone.0238616

65. Zhang, X., Liu, Y., Yuan, G., Wang, D., Zhu, T., Wu, X., Ma, M., Guo, L., Guo, H., Bhadauria, V., Liu, J., Peng, Y.-L., 2022. The effector recognition by synthetic sensor NLR receptors requires the concerted action of multiple interfaces within and outside the integrated domain. https://doi.org/10.1101/2022.08.17.504349

66. Zhang, Z.-M., Ma, K.-W., Gao, L., Hu, Z., Schwizer, S., Ma, W., Song, J., 2017. Mechanism of host substrate acetylation by a YopJ family effector. Nat Plants 3, 17115. https://doi.org/10.1038/nplants.2017.115

## References

68. De la Concepcion, J.C., Franceschetti, M., MacLean, D., Terauchi, R., Kamoun, S., Banfield, M.J., 2019. Protein engineering expands the effector recognition profile of a rice NLR immune receptor. eLife 8, e47713. https://doi.org/10.7554/eLife.47713

69. Ho, J., Tumkaya, T., Aryal, S., Choi, H., Claridge-Chang, A., 2019. Moving beyond P values: data analysis with estimation graphics. Nat Methods 16, 565–566. https://doi.org/10.1038/s41592-019-0470-3

70. MacLean, D., 2019. TeamMacLean/besthr: Initial Release. Zenodo. https://doi.org/10.5281/zenodo.3374507

71. Maqbool, A., Saitoh, H., Franceschetti, M., Stevenson, C., Uemura, A., Kanzaki, H., Kamoun, S., Terauchi, R., Banfield, M., 2015. Structural basis of pathogen recognition by an integrated HMA domain in a plant NLR immune receptor. eLife 4, e08709. https://doi.org/10.7554/eLife.08709

